# SARS-CoV-2 hijacks neutralizing dimeric IgA for enhanced nasal infection and injury

**DOI:** 10.1101/2021.10.05.463282

**Authors:** Biao Zhou, Runhong Zhou, Jasper Fuk-Woo Chan, Jianwei Zeng, Qi Zhang, Shuofeng Yuan, Li Liu, Rémy Robinot, Sisi Shan, Jiwan Ge, Hugo Yat-Hei Kwong, Dongyan Zhou, Haoran Xu, Chris Chung-Sing Chan, Vincent Kwok-Man Poon, Hin Chu, Ming Yue, Ka-Yi Kwan, Chun-Yin Chan, Na Liu, Chris Chun-Yiu Chan, Kenn Ka-Heng Chik, Zhenglong Du, Ka-Kit Au, Haode Huang, Hiu-On Man, Jianli Cao, Cun Li, Ziyi Wang, Jie Zhou, Youqiang Song, Man-Lung Yeung, Kelvin Kai-Wang To, David D. Ho, Lisa A. Chakrabarti, Xinquan Wang, Linqi Zhang, Kwok-Yung Yuen, Zhiwei Chen

## Abstract

Robust severe acute respiratory syndrome coronavirus-2 (SARS-CoV-2) infection in nasal turbinate (NT) accounts for high viral transmissibility, yet whether neutralizing IgA antibodies can control it remains unknown. Here, we evaluated receptor binding domain (RBD)-specific monomeric B8-mIgA1 and B8-mIgA2, and dimeric B8-dIgA1 and B8-dIgA2 against intranasal SARS-CoV-2 challenge in Syrian hamsters. These antibodies exhibited comparably potent neutralization against authentic virus by competing with human angiotensin converting enzyme-2 (ACE2) receptor for RBD binding. While reducing viruses in lungs, pre-exposure intranasal B8-dIgA1 or B8-dIgA2 led to 81-fold more infectious viruses and severer damage in NT than placebo. Virus-bound B8-dIgA1 and B8-dIgA2 could engage CD209 as an alternative receptor for entry into ACE2-negative cells and allowed viral cell-to-cell transmission. Cryo-EM revealed B8 as a class II neutralizing antibody binding trimeric RBDs in 3-up or 2-up/1-down conformation. Therefore, RBD-specific neutralizing dIgA engages an unexpected action for enhanced SARS-CoV-2 nasal infection and injury in Syrian hamsters.

## INTRODUCTION

Severe acute respiratory syndrome coronavirus 2 (SARS-CoV-2), a member of the *Betacoronavirus* genus, is the causative agent of Coronavirus Disease 2019 (COVID-19) ^1^. SARS-CoV-2 enters host cells through the binding of the receptor binding domain (RBD) of its surface trimeric spike (S) protein to the cellular angiotensin-converting enzyme-2 (ACE-2) receptor ^2–4^. After the ACE2 binding, the S protein is cleaved into S1 and S2 subunits by host cellular proteases including the transmembrane protease serine 2 (TMPRSS2) to promote fusion of viral and cellular membranes for viral entry ^5–8^. Apparently, these entry processes are similar to those of SARS-CoV-1 ^9–12^, the causative agent of SARS, although these two coronaviruses share just 76% and 40% amino acid identity between their genomes and their RBD external subdomains, respectively ^13,14^. SARS-CoV-2, however, has displayed remarkably higher transmissibility than SARS-CoV-1. By August 2021, the rapid all-year-round transmission of SARS-CoV-2 has resulted in over 200 million infections and 4 million deaths globally ^15^ since the outbreak of COVID-19 reported in December 2019. This is in marked contrast to the SARS epidemic, which caused only 8096 cases and 774 deaths, and disappeared in 2003 ^16^. Therefore, understanding the mechanisms underlying the high transmissibility of SARS-CoV-2 is essential for pandemic control.

The high transmissibility of SARS-CoV-2 is likely associated with multiple factors. First, unlike SARS-CoV-1-infected cases characterized by high fever and prominent respiratory symptoms, afebrile individuals with SARS-CoV-2 were often found upon diagnosis, allowing person-to-person transmissions by asymptomatic carriers including international travellers ^17–20^. Second, while both coronaviruses employ the same receptor ACE2, highly conserved RBD residues or side chain properties of SARS-CoV-2 might account for increased ACE2 binding ^9–12^. Third, the unique insertion of the PRRA sequence in SARS-CoV-2 S glycoprotein promotes higher virion infectivity and cell-cell fusion, leading to enhanced pathogenicity *in vivo* ^21–23^. Fourth, by altering the S protein conformation, the D614G mutation increases the stability of S trimer to avoid premature S1 shedding, which results in a rapid dominance of this mutation globally ^24,25^. The D614G mutation also induces higher infectious titres in nasal washes and the trachea of infected hamsters ^26^. Multiple mechanisms, therefore, contribute to the high upper respiratory tract (URT) viral loads characteristic of SARS-CoV-2 infection ^27–29^, which facilitates viral transmission in the human population. Till now, the role of nasal IgA remains understudied. In particular, whether antibody-dependent enhancement (ADE) of SARS-CoV-2 infection has any place *in vivo* remains an open question ^30^.

Since the outbreak of COVID-19, worldwide research efforts have led to the identification of many potent human neutralizing antibodies (HuNAbs) mainly in IgG form for preclinical and clinical developments ^31–38^. Some studies also investigated IgA antibodies, which are known to play an important role in mucosal immunity, especially in their secretory form (SIgA) ^39,40^. RBD-specific IgA antibodies were rapidly discovered in COVID-19 patients ^41^. IgA and SIgA were even shown to dominate the early antibody response as compared to IgG and IgM in saliva and bronchoalveolar lavage fluids due to expansion of IgA plasmablasts with mucosal homing characteristics ^42,43^. However, it was noted that COVID-19 patients with acute respiratory distress syndrome (ARDS) had higher SIgA in the airway mucosa for unknown reasons ^44^. Moreover, a recent study reported that SARS-CoV-2 viral loads were closely associated with spike-specific IgA responses in the nasal samples of acute COVID-19 patients ^45^. Since dIgA were shown to be about 15 times more potent than mIgA *in vitro* against the same target, dIgA were suggested to be particularly valuable for therapeutic application against SARS-CoV-2 ^46^. It is essential to investigate the potential of mIgA and dIgA in preventing SARS-CoV-2 infection *in vivo*.

In this study, we used the technology of single B cell antibody gene cloning to generate a panel of SARS-CoV-2 RBD-specific monoclonal HuNAbs from the peripheral blood mononuclear cells (PBMCs) of one acute and three convalescent COVID-19 patients in Hong Kong. Since intramuscular or intranasal inoculation of several potent IgG HuNAbs cannot completely prevent SARS-CoV-2 infection in the nasal turbinate (NT) of Syrian hamsters ^47^, we sought to improve the efficacy of HuNAb by converting IgG to IgA. To achieve this goal, we engineered the potent B8-IgG1 into monomeric IgA1 (B8-mIgA1), monomeric IgA2 (B8-mIgA2), dimeric (B8-dIgA1) and dimeric IgA2 (B8-dIgA2), and determined their efficacies in the Syrian hamster model against the live intranasal SARS-CoV-2 challenge ^48^.

## RESULTS

### Characterization of human monoclonal antibodies from COVID-19 patients

To isolate human monoclonal antibodies (MAbs), we obtained peripheral blood mononuclear cells (PBMCs) from one acute (P4) and three convalescent (P1-P3) COVID-19 patients in Hong Kong at a mean 73.5 (±25) days after symptoms onset (Supplementary Table 1). Enzyme-linked immunosorbent assay (ELISA) and pseudovirus neutralization assays revealed that all patient sera showed SARS-CoV-2 RBD- and spike-specific binding (Fig. 1A and 1B) and neutralizing antibody (NAb) activities (Fig. 1C). The mean NAb IC_50_ titer was 1:1753 with a range of 1:638-1:5701. Flow cytometry was then used to sort SARS-CoV-2-specific immunoglobulin positive (IgG^+^) memory B cells from individual PBMC samples using two fluorescent-conjugated RBD probes. The percentage of RBD-binding IgG^+^ memory B cells ranged from 0.19% to 0.52% (Supplementary Fig. 1A and Supplementary Fig. 1B). We successfully cloned a total of 34 MAbs from these patients, including 3 from P1, 8 from P2, 17 from P3 and 6 from P4. We confirmed that 18 of these MAbs exhibited RBD-specific binding activities detected by ELISA (Supplementary Fig. 1C). No clear dominance of heavy (H) chain gene family was found among these 4 subjects by sequence analysis (Fig. 1D, left panels). VLK1, however, was the most used variable gene family for the light (L) chain (Fig. 1D, right panels). The average somatic hypermutation (SHM) rate ranged from 0% to 12.2% for the H chain and from 0.7% to 7.9% for the L chain (Fig. 1E, left). The average complementarity-determining region 3 (CDR3) lengths ranged from 12.3 to 17.4 for the H chain and 8.4 to 9.4 for the L chain, respectively (Fig. 1E, right). These results suggested overall comparable degrees of affinity maturation in these RBD-specific human MAbs obtained from individual memory B cells of four Hong Kong patients.

**Fig. 1.**
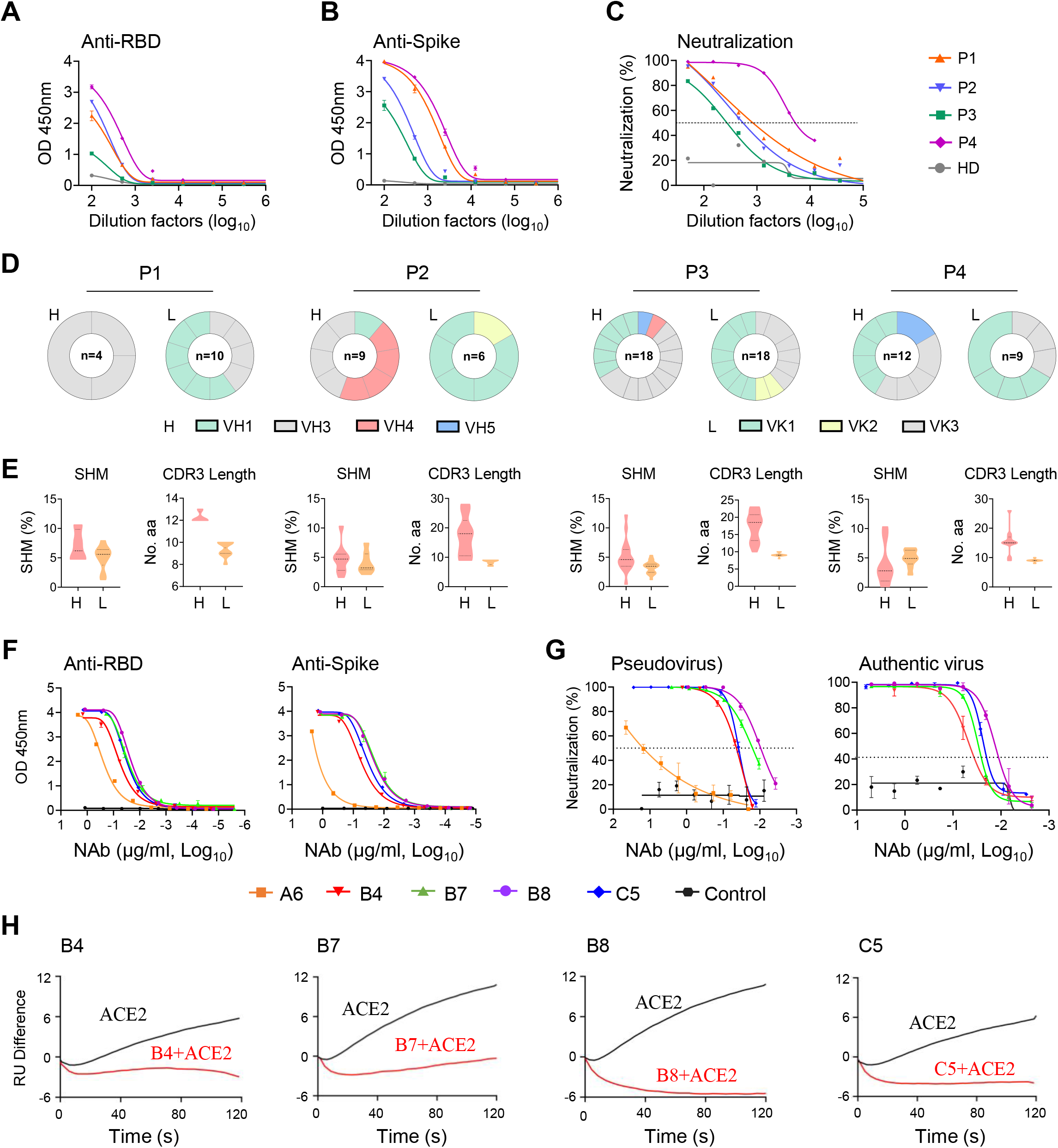
Isolation of monoclonal antibodies from single B cells of convalescent COVID-19 patients. **(A)** RBD-specific binding activities of sera derived from 3 (P1-P3) convalescent and 1 (P4) acute COVID-19 patients as measured by ELISA. **(B)** Spike-specific binding activities of sera derived from four COVID-19 patients as measured by ELISA. **(C)** Neutralization activities of sera derived from four COVID-19 patients as measured by pseudotyped SARS-CoV-2 inhibition in 293T-ACE2 cells. **(D)** Antibody gene repertoire analysis of reactive B cells derived from each patient. The number of cloned antibody genes from each patient is shown in the center of each pie chart for both the heavy (H) and light (L) chains. The colors represent specific variable gene family. Each fragment of the same color stands for one specific sub-family. **(E)**. The percentage of somatic hypermutation (SHM) compared to germline sequences and the CDR3 amino acid lengths of cloned antibody H and L gene sequences were analyzed for each subject. **(F)** RBD (left) and spike (right) specific binding activities of five HuNAbs, including A6, B4, B7, B8 and C5, were measured by ELISA. **(G)** Neutralization activities of 5 HuNAbs against pseudotyped (left) and authentic (right) SARS-CoV-2 were determined in HEK 293T-ACE2 and Vero-E6 cells, respectively. HIV-1 specific HuNAb VRC01 served as a negative control. Each assay was performed in duplicates and the mean of replicates is shown with the standard error of mean (SEM).s **(H)** The competition of four HuNAbs, including B4, B7, B8 and C5, with human soluble ACE2 for binding to SARS-CoV-2 RBD was measured by SPR. The curves show binding of ACE2 to SARS-CoV-2 RBD with (red) or without (black) pre-incubation with each HuNAb.

### Specificity and potency of SARS-CoV-2-specific human neutralizing antibodies

To determine the antiviral activities of these 18 RBD-specific human MAbs, we performed binding and neutralization assays. Five of them, namely A6-IgG1, B4-IgG1, B7-IgG1, B8-IgG1 and C5-IgG1 displayed RBD- and spike- specific binding by ELISA (Fig. 1F) and neutralizing activities against both pseudotyped and authentic viruses (Fig. 1G and Supplementary Fig. 1D). Interestingly, the four most potent HuNAbs, B4-IgG1, B7-IgG1, B8-IgG1 and C5-IgG1, all came from patient P3 (Supplementary Fig. 1D). Sequence analysis revealed strong similarities between B7-, B8-, and C5-IgG1, which all contained an IGHV1-69 heavy chain gene and an IGKV3 kappa light chain gene. B4-IgG1 contained distinct IGHV3-66 and IGKV1-33 genes with CDR3 lengths of 12 amino acids (aa) and 9 aa, and somatic hypermutation (SHM) rate of 3.8% and 4.6%, respectively (Supplementary Table 2). B7 and B8 were the most similar, as both contained IGHV1-69 and IGKV3-11 though B8 had a shorter CDR3 (14 aa vs 18 aa) and higher SHM (4.8% vs 0.0%) than those of B7 in the heavy chain. In the L chain, B7 and B8 shared a 9 aa CDR3, but B8 had a higher SHM rate than that of B7 (1.7% vs 0.7%) C5-IgG1 had a similar IGHV1-69 light chain gene with a 16 aa CDR3 and 2.4% SHM, but a different IGKV3-20 with a 9 aa CDR3 and 2.8% SHM. By ELISA, these four P3-derived HuNAbs bound to the SARS-CoV-2 RBD with half-maximal effective concentration (EC_50_) values ranging from 0.02 to 0.06 μg/ml, indicating stronger binding than that of the P4-derived A6-IgG1 (0.3 μg/ml) (Fig. 1F, left and Supplementary Table 3). P3-derived MAbs also exhibited stronger binding activities to the spike, with the EC_50_ values ranging from 0.018 to 0.06 μg/ml, compared to A6-IgG1 (17.94 μg/ml) (Fig. 1F, right and Supplementary Table 3). Neutralizing assays using pseudoviruses revealed that these four potent HuNAbs had IC_50_ values ranging from 0.0095 to 0.038 μg/ml, and IC_90_ values ranging from 0.046 to 0.136 μg/ml, respectively (Fig. 1G, left and Supplementary Table 4). Moreover, B8 proved to be the most potent HuNAb, capable of inhibiting authentic SARS-CoV-2 with an IC_50_ value of 0.013 μg/ml and an IC_90_ value of 0.032 μg/ml, respectively (Fig. 1G, right and Supplementary Table 4). We then determined if these HuNAbs could compete with ACE2 for RBD binding by surface plasmon resonance (SPR). We found that all of them strongly competed with ACE2 (Fig. 1H). In line with the ELISA results, B8-IgG1 displayed the best KD value for RBD binding (169 pM) (Supplementary Fig. 2A, Supplementary Table 5) and the strongest competition with ACE2 (Fig. 1H), which explained its potent neutralizing activity. As B4-IgG1 displayed only partial competition for RBD binding with the other antibodies (Supplementary Fig. 2B), we further performed antibody synergy experiments using the pseudotype neuralization assay. No significant synergistic effects were found between any pairs of these four HuNAbs (Supplementary Fig. 2C). These results demonstrated that B4-, B7-, B8- and C5-IgG1 were all RBD-specific and competed with ACE2 for similar sites on the RBD.

### B8-IgG1 pre-exposure prophylaxis and post-exposure treatment in the golden Syrian hamster model

To determine the efficiency of B8-IgG1 in pre-exposure prophylaxis and post-exposure treatment against live intranasal SARS-CoV-2 infection, we administered B8-IgG1 intraperitoneally in golden Syrian hamsters, before or after viral challenge in our Biosafety Level-3 (BSL-3) animal laboratory (Fig. 2A). In the pre-exposure prophylaxis group (G1, n = 4), each hamster received a single intraperitoneal injection of 1.5 mg/kg B8-IgG1. In the post-exposure treatment groups, each hamster received a single intraperitoneal injection of 1.5 mg/kg B8-IgG1 at day 1 (G2, n = 4), day 2 (G3, n = 4) or day 3 (G4, n = 4) postinfection (dpi), respectively (Fig. 2A). The challenge dose was 10^5^ plaque-forming units (PFU) of live SARS-CoV-2 (HKU-001a strain) ^47,48^. Another group of hamsters (G0, n = 4) received PBS injection as a no-treatment control. One G4 animal died accidentally during the procedure. Since Syrian hamsters recover quickly from SARS-CoV-2 infection, with resolution of clinical signs and clearance of virus shedding within one week after infection ^26,48^, we chose to sacrifice the animals at 4 dpi for HuNAb efficacy analysis, at a time when high viral loads and acute lung injury were consistently observed. At 4 dpi, NT and lung tissues were harvested to quantify infectious viruses by measuring PFUs, viral RNA loads by real-time reverse-transcription polymerase chain reaction (RT-PCR) and infected cells by immunofluorescence (IF) staining of viral nucleocapsid protein (NP)-positive cells as we described previously ^47,48^.

**Fig. 2.**
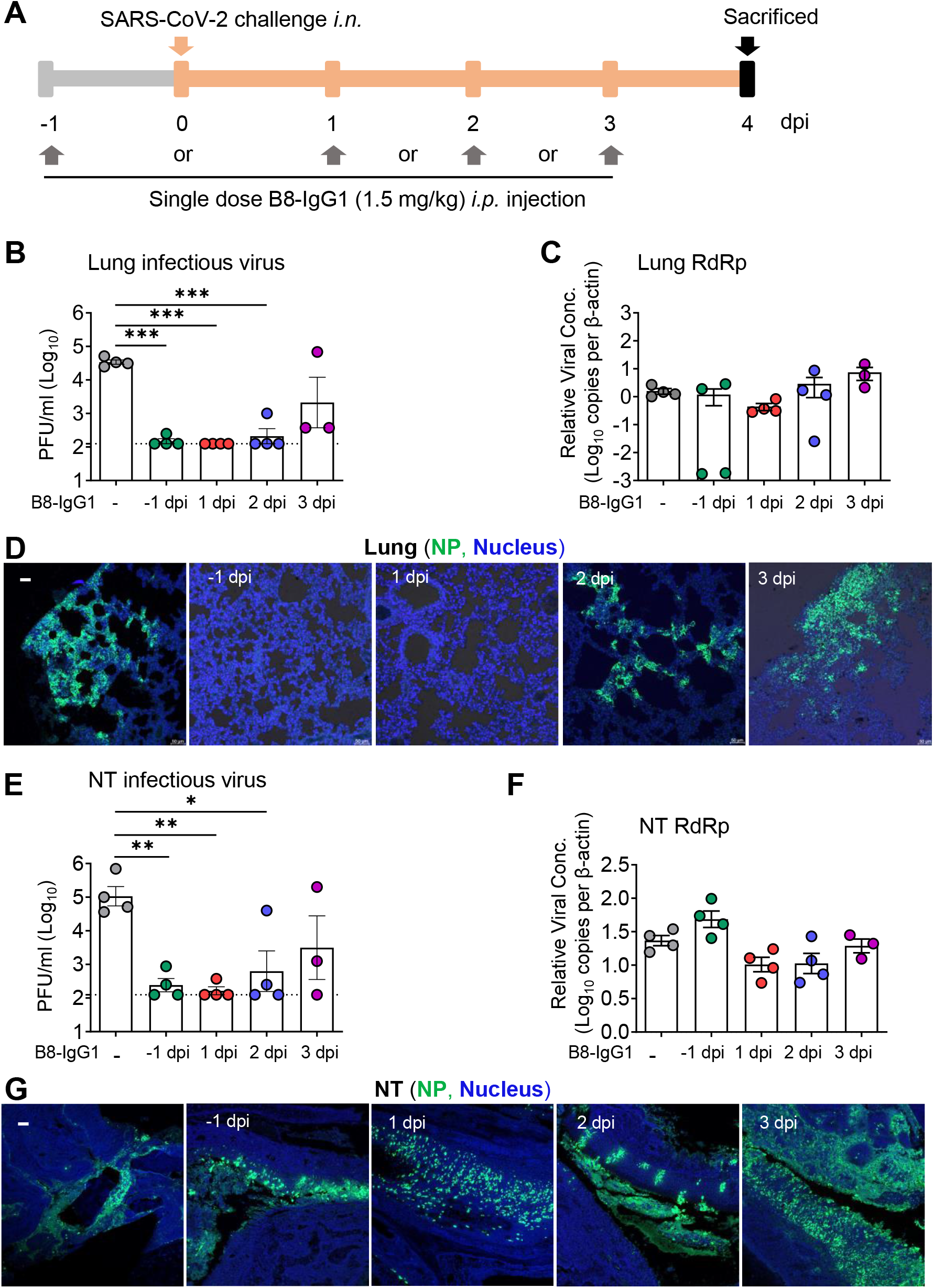
Pre- and post-exposure treatment of B8-IgG1 against SARS-CoV-2 in Syrian Hamster. **(A)** Experimental schedule. Four groups of hamsters (G1-G4) received intraperitoneally a single dose of 1.5 mg/kg of B8-IgG1 at one day before infection (−1 dpi) for pre-exposure prophylaxis, and at day one (1 dpi), two (2 dpi) and three (3 dpi) post-infection for early treatment, respectively. Control hamsters (G0, n=4) received an isotypic control antibody at the same dose. On day 0, each hamster was intranasally challenged with a dose of 10^5^ PFU of SARS-CoV-2 (HKU-001a strain). All hamsters were sacrificed on 4 dpi for analysis. **(B)** Infectious virus (PFU) was measured in animal lungs by the viral plaque assay in Vero-E6 cells. The PFU/ml concentration is shown in log-transformed units. **(C)** The relative viral RdRp RNA copies (normalized to β-actin) were determined by RT-PCR in animal lungs. **(D)** Representative 100× images of infected lungs from each group, as determined by anti-NP immunofluorescence (IF) staining. The cell nuclei were counterstained with DAPI (blue). **(E)** Infectious virus (PFU) were measured in NT homogenates by the viral plaque assay as mentioned above. **(F)** Viral loads in NT homogenates of each group were determined by RT-PCR assay. The viral load data is shown in log-transformed units. **(G)** Representative 100× images of infected NT from each group as determined by anti-NP IF staining as mentioned above. Statistics were generated using one-way ANOVA tests. \**p*<0.05; \*\**p*<0.01; \*\*\**p*<0.001.

We found that infectious virus, measured by PFU, weas readily detected in all tissue compartments of G0 hamsters but not in the lungs of 75% G1, 100% G2, 75% G3 and 0% G4 animals, nor in the NT of 50% G1, 75% G2, 50% G3 and 25% G4 animals (Fig. 2B and 2E). The decrease in PFU was of 2-3 orders of magnitude, suggesting efficient viral suppression in the lungs for the G1, G2, and G3 groups. A sensitive RT-PCR assay further demonstrated that viral RNA copy numbers were decreased in the lungs by 3 orders of magnitude in 50% of G1 hamsters (Fig. 2C). In contrast, there was no significant viral RNA reduction in the NT of G1 animals (Fig. 2F), suggesting lower efficacy of B8-IgG1 to prevent viral entry in the URT than in the lungs. There were slight but not significant viral RNA load decreases in both lungs and NT of G2 and some G3 animals. We then evaluated the number of infected cells or foci in these two tissue compartments by anti-NP antibody staining. A clear decrease of NP-positive cells or foci was observed in the lungs of G1 and G2 hamsters (Fig. 2D). Abundant NP-positive cells with a diffuse distribution, however, were readily detected in the NT of all the challenged hamsters (Fig. 2G). These results demonstrated that systemic B8-IgG1 injection was effective at reducing productive SARS-CoV-2 infection in the lungs when used for pre-exposure prophylaxis and early treatment especially within 48 hours post infection, but was insufficient to prevent viral infection in the NT.

To determine the correlates of B8-IgG1-mediated protection, we also measured the antibody concentration in serum, lung homogenate and NT homogenates at 0 and 4 dpi for all experimental animals. On average, 4,257 ng/ml and 2,101 ng/ml B8-IgG1 were found in animal sera at 0 and 4 dpi (Supplementary Table 6). On 4 dpi, lung and NT homogenates contained 128 ng/ml and 20 ng/ml in G1, 238 ng/ml and 86 ng/ml in G2, 229 ng/ml and 93 ng/ml in G3, and 192 ng/ml and 46 ng/ml in G4 animals, respectively. These results demonstrated that most animals maintained higher peripheral B8-IgG1 antibody concentration, while a decreasing concentration gradient was observed in the lungs and NT during infection. The concentrations of B8-IgG1 measured in NT homogenates were close to the neutralization IC_90_ measured *in vitro* (32 ng/ml), explaining why infectious virus was undetectable in the PFU assay. These findings are in line with results obtained for other potent IgG HuNAbs administered systemically, as reported in our previous study ^47^. The B8-IgG1 concentrations measured in the NT appeared insufficient to completely block infection *in vivo*, as indicated by the presence of NP-positive cells scattered throughout the NT.

### Pre-exposure prophylaxis by monomeric B8-mIgA1 and B8-mIgA2 in Syrian hamsters

Since systemic administration of the RBD-specific neutralizing B8-IgG1 did not suppress SARS-CoV-2 nasal infection effectively, we sought to construct various types of IgA for mucosal intervention. For this purpose, we engineered B8-IgG1 into monomeric B8-mIgA1 and B8-mIgA2, and then into dimeric B8-dIgA1 and B8-dIgA2 by introducing the J chain. By *in vitro* characterization, purified B8-mIgA1, B8-mIgA2, B8-dIgA1 and B8-dIgA2 retained similar binding to RBD and spike by ELISA as compared to B8-IgG1 (Supplementary Fig. 3A-B, Supplementary Table 7), and comparable antiviral activities based on neutralization IC_50_ and IC_90_ values ((Supplementary Fig. 3C-D and Supplementary Table 8). That said, B8-mIgA2 and B8-dIgA2 showed slightly more potent IC_90_ activities than B8-IgG1 in the pseudovirus neutralization assay using 293T-ACE2 cells as targets (Supplementary Table 8). After introducing J chain, the proper dimer formation of B8-dIgA1 and B8-dIgA2 was confirmed by size exclusion chromatography analysis (Supplementary Fig. 3E-F). Furthermore, B8-mIgA1, B8-mIgA2, B8-dIgA1 and B8-dIgA2 also retained comparable competition with ACE2 for binding to spike by SPR analysis (Supplementary Fig. 3G-J). Therefore, the engineered IgA had the expected structural properties, and showed antiviral activities as potent as those of B8-IgG1 *in vitro*.

We then evaluated the monomeric B8-mIgA1 and B8-mIgA2 in the hamster model, using a higher 4.5 mg/kg dose via either intranasal or intraperitoneal injection, and used the same amount of intranasal B8-IgG1 inoculation as a control (Fig. 3A to 3I). Interestingly, while changes in total RdRp and subgenomic sgNP viral RNA loads were not obvious (Fig. 3D to 3E), B8-IgG1 and B8-mIgA1 (both *i.n.* and *i.p.*), but not B8-mIgA2, were able to significantly suppress infectious virus production (PFU) in the lungs of 75% infected hamsters by 2 orders of magnitude (Fig. 3D). Sporadic infected cell foci were still detected in the lung sections by anti-NP staining (Fig. 4E and Supplementary Fig. 4A), suggesting that protection conferred by B8-IgG1 and B8-mIgA1 was not complete. B8-mIgA2 was not able to suppress viral RNA load nor PFU in the challenged hamsters, regardless of the route of antibody injection. On the other hand, like B8-IgG1, both B8-mIgA1 and B8-mIgA2 did not achieve significant viral suppression in the NT. After intranasal administration of either B8-mIgA1 or B8-mIgA2, some hamsters even showed a trend of slightly increased infectious virus production in NT, though this did not reach statistical significance (Fig. 3F-3H). Among these animals, the NP-positive cells were detected readily in the NT, as demonstrated by the whole section scanning, indicating the comparable distribution compared with the B8-IgG1 and no-treatment groups (Fig. 3I). These results were consistent with many NP-positive cells observed in diffusely infected areas of NT by classic IF (Supplementary Fig. 4B). These results demonstrated that B8-mIgA1 was more potent than B8-mIgA2 at limiting SARS-CoV-2 infection in the lungs, but that both mIgA did not prevent nor significantly limit SARS-CoV-2 nasal infection.

**Fig. 3.**
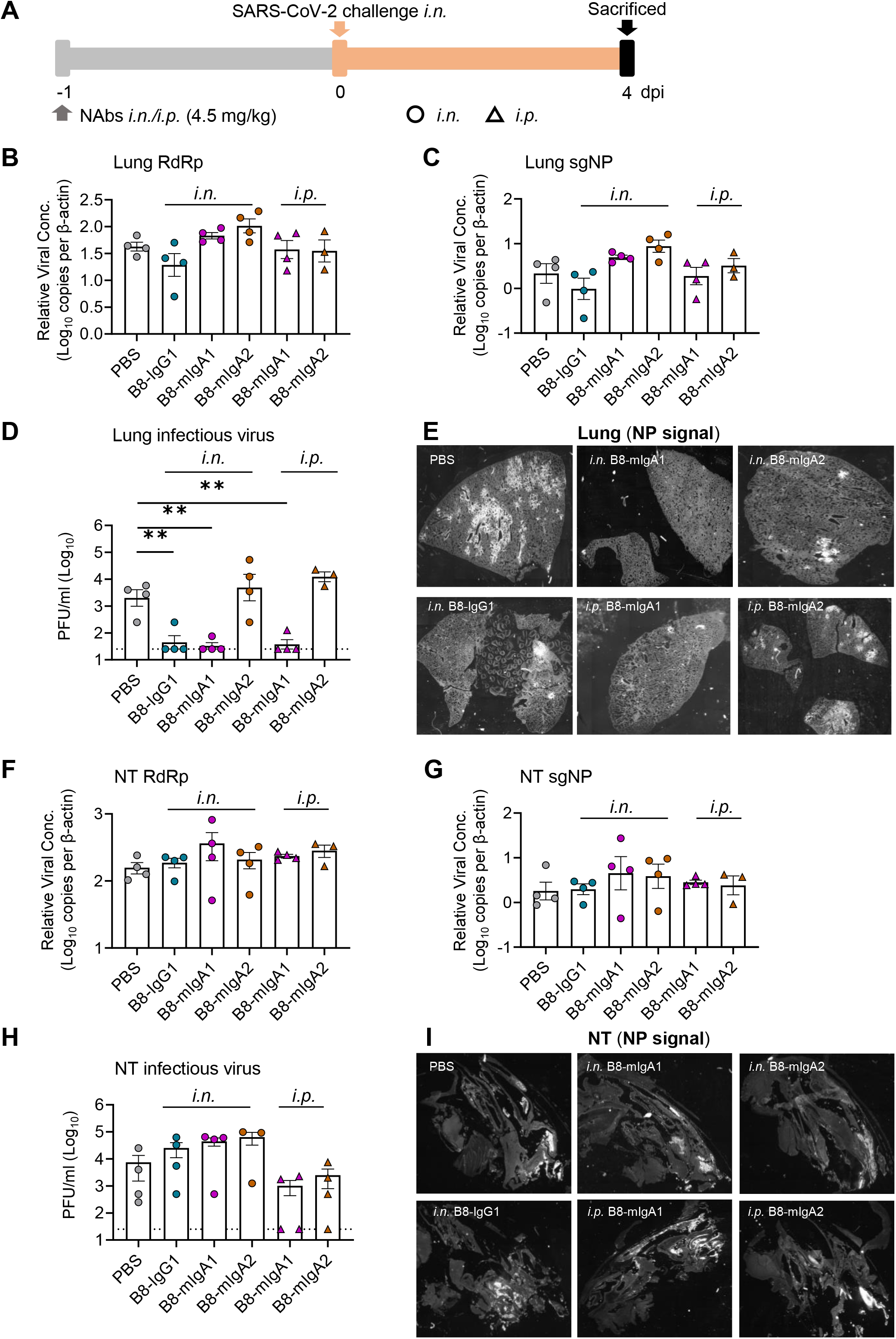
Pre-exposure treatment with monomeric B8-mIgA1 or B8-mIgA2 against SARS-CoV-2 infection in Syrian hamsters. **(A)** Experimental schedule. Five groups of hamsters (n=4 per group) received a single dose of 4.5 mg/kg of B8-IgG1, B8-mIgA1 or B8-mIgA2 one day before viral challenge for pre-exposure prophylaxis by the intranasal route (circles) or the intraperitoneal route (triangles), respectively. Control hamsters (n=4) received PBS. On day 0, each hamster was intranasally challenged with a dose of 10^5^ PFU of SARS-CoV-2, as mentioned in Fig. 2A. All hamsters were sacrificed on 4 dpi for analysis. **(B)** The viral RNA load, measured by relative RdRp RNA copy numbers (normalized to β-actin) was determined by RT-PCR in animal lung homogenates. **(C)** The relative sub-genomic nucleocapsid (sgNP) RNA copy numbers (normalized to β-actin) were determined by RT-PCR in animal lung homogenates. **(D)** Infectious virus (PFU) was measured in animal lung homogenates by the viral plaque assay in Vero-E6 cells. **(E)** Representative lung images of infected animals by scanning whole tissue section. The signal of SARS-CoV-2 NP was shown in bright spots. **(F)** The relative viral RdRp RNA copy numbers (normalized to β-actin) were determined by RT-PCR in NT homogenates. **(G)** The relative sgNP RNA copy numbers (normalized to β-actin) were determined by RT-PCR in NT homogenates. **(H)** Infectious virus (PFU) was measured in animal NT homogenates by the viral plaque assay in Vero-E6 cells. **(I)** Representative NT images of infected animals by scanning whole tissue section. The signal of SARS-CoV-2 NP was shown in bright spots. Log-transformed units are shown in **(B) to (H)** except in **(E)**. Statistics were generated using one-way ANOVA tests. \**p*<0.05; \*\**p*<0.01.

**Fig. 4.**
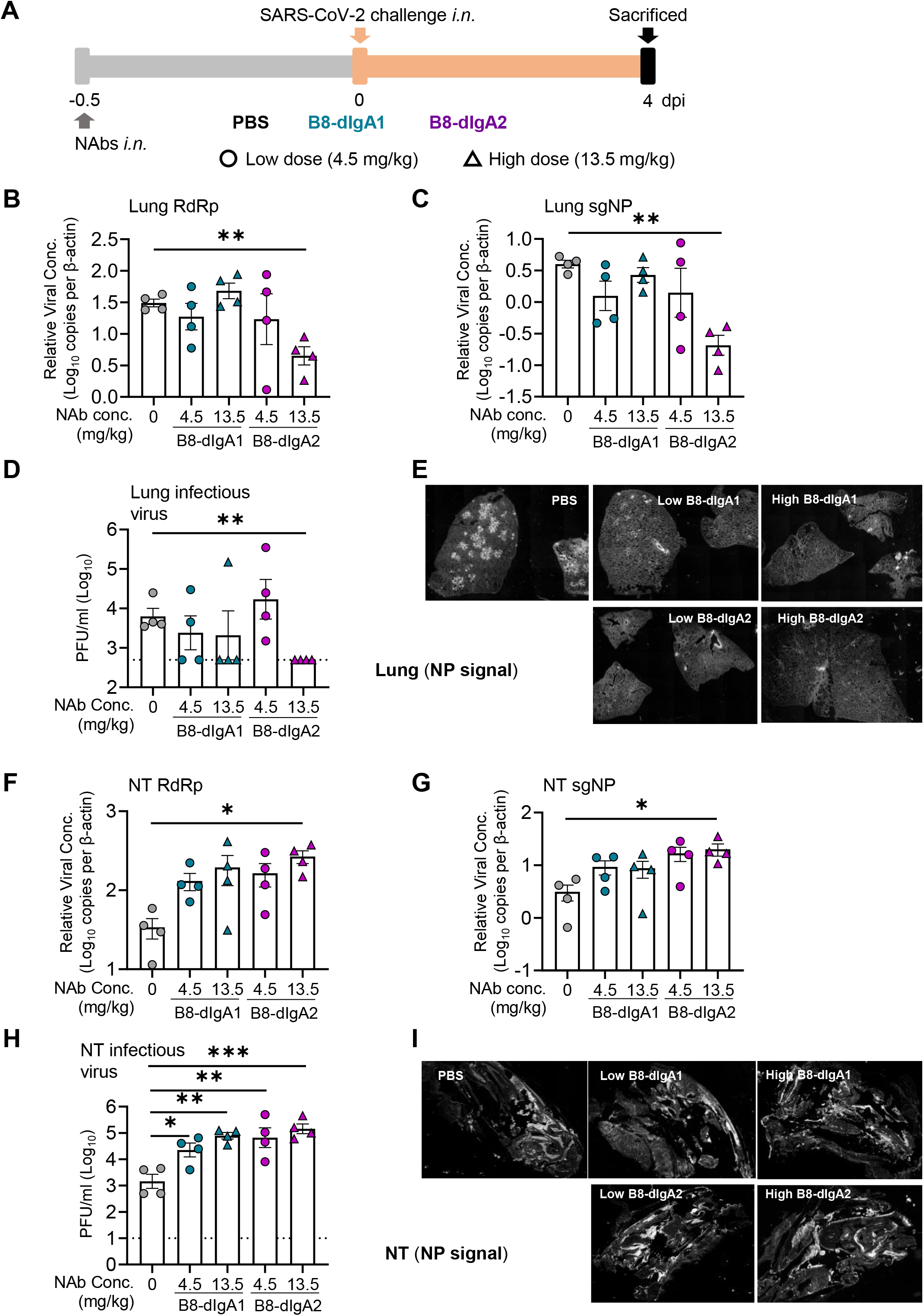
Pre-exposure treatment with dimeric B8-dIgA enhances SARS-CoV-2 infection in Syrian hamsters. **(A)** Experimental schedule. Four groups of hamsters (n=4 per group) were inoculated intranasally with B8-dIgA1 or B8-dIgA2, either at a low dose of 4.5 mg/kg or at a high dose of 13.5 mg/kg, respectively, 12 hours before intranasal viral challenge. Another group of hamsters (n=4) received PBS as control. On day 0, each hamster was intranasally challenged with a dose of 10^5^ PFU of SARS-CoV-2 as described in Fig. 2A. All hamsters were sacrificed on 4 dpi for analysis. Data represent a presentative experiment from three independent experiments. **(B)** The relative viral RdRp RNA copy numbers (normalized to β-actin) were determined by RT-PCR in animal lung homogenates. **(C)** The relative viral sgNP RNA copy numbers (normalized to β-actin) were determined by RT-PCR in animal lung homogenates. **(D)** Infectious virus (PFU) was measured in animal lung homogenates by the viral plaque assay in Vero-E6 cells. **(E)** Representative lung images of infected animals by scanning whole tissue section. The signal of SARS-CoV-2 NP was shown in bright spots. **(F)** The relative viral RdRp RNA copy numbers (normalized to β-actin) were determined by RT-PCR in NT homogenates. **(G)** The relative viral sgNP RNA copies (normalized to β-actin) were determined by RT-PCR in NT homogenates. **(H)** Infectious virus (PFU) was measured in NT homogenates by the viral plaque assay in Vero-E6 cells. **(I)** Representative NT images of infected animals by scanning whole tissue section. The signal of SARS-CoV-2 NP was shown in bright spots. Log-transformed units are shown in **(B) to (H)** except in **(E)**. Statistics were generated using one-way ANOVA tests. \**p*<0.05; \*\**p*<0.01.

### B8-dIgA1 and B8-dIgA2 mediate enhancement of SARS-CoV-2 nasal infection and injury in Syrian hamsters

Next, we tested the effects of dimeric B8-dIgA1 and B8-dIgA2 in Syrian hamsters. To improve protective efficacy of intranasal dIgA treatment, we included a 3-fold higher dosage group of 13.5 mg/kg besides the 4.5 mg/kg group (Fig. 4A to 4I), and we shortened the interval between dIgA and virus inoculation to 12 hours (Fig. 4A). Both RdRp and sgNP viral RNA loads dropped significantly in the lungs of hamsters that received the high dose of B8-dIgA2 compared to the no-treatment group (Fig. 4B and 4C). Both B8-dIgA1 and B8-dIgA2 at the high dose also suppressed infectious viruses (PFU) in the lungs of 75% and 100% treated hamsters, respectively (Fig. 4D). High dose B8-dIgA1 and B8-dIgA2 also decreased the number of NP-positive cells or foci in the lungs, with a more marked change for B8-dIgA2 (Fig. 4E and Supplementary Fig. 5A). Unexpectedly, however, we observed significantly enhanced SARS-CoV-2 nasal infection and tissue damage in most infected hamsters included the low and high dose groups of both B8-dIgA1 and B8-dIgA2, in all the four assays used (Fig. 4F to 4I). High dose administration of B8-dIgA1 or B8-dIgA2 resulted in increased PFU production in the NT by 37-fold and 81-fold, respectively, compared to the no-treatment group (Fig. 4H). Since our model showed comparable NT PFU on day 2 and day 4 as described previously ^48^, this level of enhanced infection was unusual. It was also not observed with B8-IgG1 or monomeric B8-mIgA1 and B8-mIgA2 treatment, as described above. Moreover, the distribution of NP-positive cells in hamsters treated with dimeric B8-IgA was broader and reached deeper into NT tissue compared to the no-treatment group, as shown by whole section scanning (Fig. 4I and Supplementary Fig. 5B), which was associated with more severe and extensive epithelium desquamation and luminal cell debris (Supplementary Fig. 5C). The density of nasal NP^+^ cells was also significantly higher in B8-dIgA2-treated hamsters than in PBS-treated animals (Supplementary Fig. 5D-E). It is therefore conceivable that treatment with dimeric B8-IgA expanded the type and distribution of target cells in the nasal epithelium. Critically, we confirmed that control dimeric dIgA1 and dIgA2 did not enhance SARS-CoV-2 infection under the same experimental conditions (Supplementary Fig. 6). These results demonstrated that, instead of inducing viral suppression, pre-exposure dimeric B8-dIgA1 and B8-dIgA2 enhanced SARS-CoV-2 nasal infection and injury significantly in Syrian hamsters, which was consistently found in three independent experiments.

To validate the role of B8-dIgA1 and B8-dIgA2, we then measured B8-IgA concentrations in the serum at day 0 and 4 dpi, and in the lung and NT homogenates at 4 dpi. B8-dIgA1 and B8-dIgA2 were primarily detected in lung homogenates at 4 dpi and were apparently undetectable in the serum and NT homogenates (Supplementary Table 9). The enhanced viral replication in NT probably exhausted B8-dIgA1 and B8-dIgA2 locally, through antibody-virus complex formation and clearance ^30^. To address this possibility, we treated separately five groups of naïve Syrian hamsters (n=4 per group) with each antibody at the 4.5 mg/kg dose. Twelve hours after the inoculation, antibody concentrations were readily detected in each tissues compartment (Supplementary Table 10). The highest concentrations of dIgA antibodies were found in lung homogenates, followed by nasal washes, NT homogenates, and serum. These results suggest that B8-dIgA1 and B8-dIgA2 concentrations in the NT and lungs were still above their neutralization IC_90_ values at the time of viral challenge, indicating that the results obtained in our experiments could not be explained by the insufficient amounts of antibodies.

### B8-dIgA1- and B8-dIgA2-mediated enhancement of SARS-CoV-2 infection via CD209

Since B8 antibodies share the same binding site to the RBD domain, we sought to investigate possible mechanisms of B8-dIgA1- and B8-dIgA2-enhanced infection. First, we consistently found that 10 ng/ml B8-dIgA1 or B8-dIgA2 completely neutralized SARS-CoV-2 infection in human renal proximal tubule cells (HK-2), as measured in our previously reported immunofluorescence assay (Supplementary Fig. 7) ^49^. We also tested B8-mIgA2 and B8-dIgA2 neutralizing capacity in the MucilAir™ model, consisting of a reconstructed human nasal epithelium, which contained goblet, ciliated, and basal cells (Fig. 5A) ^50^. Both B8-mIgA2 and B8-dIgA2 neutralized SARS-CoV-2 in a dose-dependent fashion, when compared to a dIgA2 control antibody. Similar experiments were carried out in the presence of the mucus naturally secreted by goblet cells, to determine whether dIgA interaction with the mucus may alter their neutralization capacity. However, B8-dIgA2 showed the same neutralization capacity in the presence and absence of mucus. These results demonstrated that B8-dIgA1 and B8-dIgA2 did not enhance SARS-CoV-2 infection in either human HK-2 or primary airway epithelial cells, which primarily expressed human ACE2 as a viral receptor. We then turned our attention to ACE2-independent mechanisms that might be associated with dimeric IgA-mediated enhancement of SARS-CoV-2 infection. Considering that mucosal monocyte-derived dendritic cells (DC) could mediate SARS-CoV-1 infection and dissemination in rhesus monkeys as early as 2 dpi, as we previously reported ^51^, we sought to investigate the role of DC-expressed surface receptors. We focused on CD209 (DC-SIGN), because this lectin was previously shown to act as a cellular receptor for secretory IgA ^52^. By IF staining, intranasal administration of B8-dIgA2 alone did not increase CD209 expression in the NT of treated hamsters (Fig. 5B, left). Upon SARS-CoV-2 infection, however, we noted an increase in CD209-positive cells in olfactory epithelium devoid of ACE2 expression (Fig. 5B, middle). Importantly, most CD209^+^ cells were positive for NP (Fig. 5B, right), indicating that these CD209^+^ cells were likely permissive to SARS-CoV-2 infection. We then determined whether B8-dIgA1 and B8-dIgA2 could enhance SARS-CoV-2 infection in 293T cells expressing human CD209 or CD299 but not ACE2. Using a low MOI of 0.05, we found that pre-incubation of B8-dIgA1 and B8-dIgA2 enhanced live SARS-CoV-2 infection significantly in 293T cells expressing human CD209, as determined by increased viral NP production (Fig. 5C). Interestingly, human CD299, a type II integral membrane protein that is 77% identical to CD209, did not show similar activities in the same experiment (Fig. 5C). Control dIgA1 and dIgA2 did not show any enhancement in NP^+^ cell detection compared with virus only. Considering that CD209^+^ DCs promote HIV-1 transmission to CD4^+^ T cells via cell-cell contacts, we speculated that B8-dIgA1 and B8-dIgA2 might not be able to block the similar process for SARS-CoV-2. Indeed, by testing the B8 antibodies at concentrations 100-times higher than IC_90_ neutralization values (around 3000 ng/ml), none of B8-IgG1, B8-mIgA1, B8-mIgA2, B8-dIgA1 and B8-dIgA2 could block cell-cell fusion (Fig. 5D). Taken together, our results demonstrated that B8-dIgA1- and B8-dIgA2-enhanced SARS-CoV-2 nasal infection likely involved viral capture and infection of mucosal CD209^+^ cells, followed by more robust infection of ACE2^+^ epithelial cells through trans-infection via cell-cell spread in NT.

**Fig. 5.**
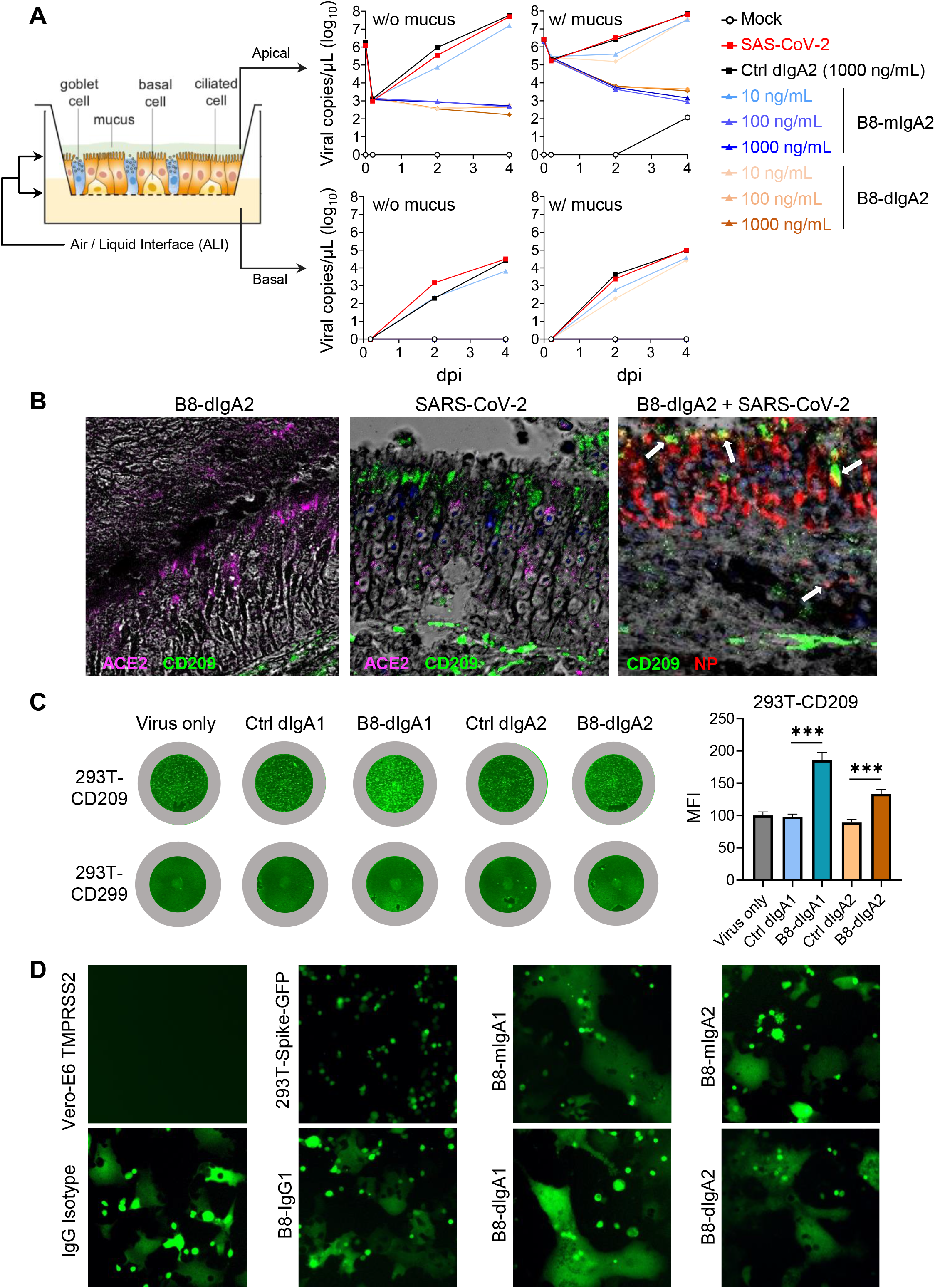
B8-dIgA1 and B8-dIgA2 enhance SARS-CoV-2 infection via CD209. **(A)** The effects of B8-dIgA2 on SARS-CoV-2 infection in the MucilAir™ model, consisting of primary human nasal epithelial cells but no DCs. B8-mIgA2 or B8-dIgA2 were pre-incubated at doses of 10, 100, and 1000 ng/ml, respectively, in the apical compartment with or without mucus for 1 hour, before adding 10^4^ PFU of SARS-CoV-2 (BetaCoV/France/IDF00372/2020) for 4 hours. The viral RNA loads were measured by RT-PCR in both the apical and basal compartments and are shown in log-transformed units. **(B)** Representative confocal images (400×) of olfactory epithelium in NT showed the expression of CD209 (DC-SIGN) in green and ACE2 in magenta by immunohistochemical staining of experimental hamsters treated with B8-dIgA2 without (left) or with (middle and right) SARS-CoV-2 infection. Color-coding indicates specific antibodies used for double staining. Infected CD209^+^ cells are visualized in yellow as indicated by arrows (right). **(C)** The CD209 or CD299 overexpressed-HEK 293T cells were pre-treated for 6 hours with 10 ng/ml of B8-dIgA1 or B8-dIgA2 or control dIgA1 or control dIgA2 or PBS, respectively, prior to SARS-CoV-2 infection (MOI: 0.05). Two days after infection, SARS-CoV-2 NP expression (green) was quantified by the mean fluorescence intensity (MFI) after anti-NP IF staining. Statistics were generated using student-*t* tests. \**p*<0.05; \*\**p*<0.01; \*\*\**p*<0.001. **(D)** The effects of B8 antibodies on cell-cell fusion. 293T cells co-transfected with SARS-CoV-2 spike and GFP were pre-treated with 100× the IC_90_ dose of B8-IgG1, B8-mIgA1, B8-mIgA2, B8-dIgA1, B8-dIgA2 or and IgG isotypic control for 1 hour, respectively. Vero-E6 cells transfected with TMPRSS2 were then added to the treated 293T-spike-GFP cells and co-cultured for 48 hours. Cell-cell fusion was imaged under a fluorescence confocal microscope at the 50× magnification.

### Cryo-EM analysis of the spike-B8 complex

To understand the potential mechanism of action of B8 HuNAb, we performed a cryo-EM single-particle analysis of B8 Fab bound to the SARS-CoV-2 spike ectodomain trimer (Supplementary Fig. 8). Two B8-spike complex structures were determined based on 351,095 and 616,799 particles collected, respectively (Supplementary Table 11). One structure at 2.67 Å resolution contained the spike with all three RBDs adopting the “up” conformation (3u), where each “up” RBD was bound by one B8 Fab (Fig. 6A-B). The other structure at 2.65 Å resolution contained one spike trimer with 2 RBDs in the “up” conformation and 1 RBD in the “down” conformation (2u1d), where each RBD was also bound by one B8 Fab despite the presence of two distinct RBD conformations (Fig. 6C-D). After superimposing the “3u” and “2u1d” spikes, a ~53-degree rotation was observed between the “up” RBD (red color) in the 3u spike trimer and the “down” RBD (gray) in the 2u1d spike trimer (Fig. 6E). The B8 Fab appeared to bind to the receptor-binding motif (RBM) of the RBD through its heavy chain for most of the interactions (Fig. 6F-G). Therefore, the cellular receptor ACE2 would clash with the B8 Fab due to the overlap of their respective epitopes on the RBM (Fig. 6F-H and Supplementary Table 12). The elucidation of the epitope revealed that B8 could be grouped into the SARS-CoV-2 neutralizing antibody class II ^53^. These structural findings were further supported by neutralization assays using a panel of pseudoviruses containing naturally occurring mutations. Indeed, the E484K mutation from the South African SA□9 strain, which is located within the B8-binding interface, caused a major loss of neutralizing potency for all the B8 isotypes tested: IgG1, mIgA1, mIgA2, dIgA1 and dIgA2 (Supplementary Table 13). The comparable neutralization profiles of these NAbs against the full panel of viral variants also indicated that the conformation of key RBD-binding residues remained unchanged after engineering of the constant regions of these B8 isotypes.

**Fig. 6.**
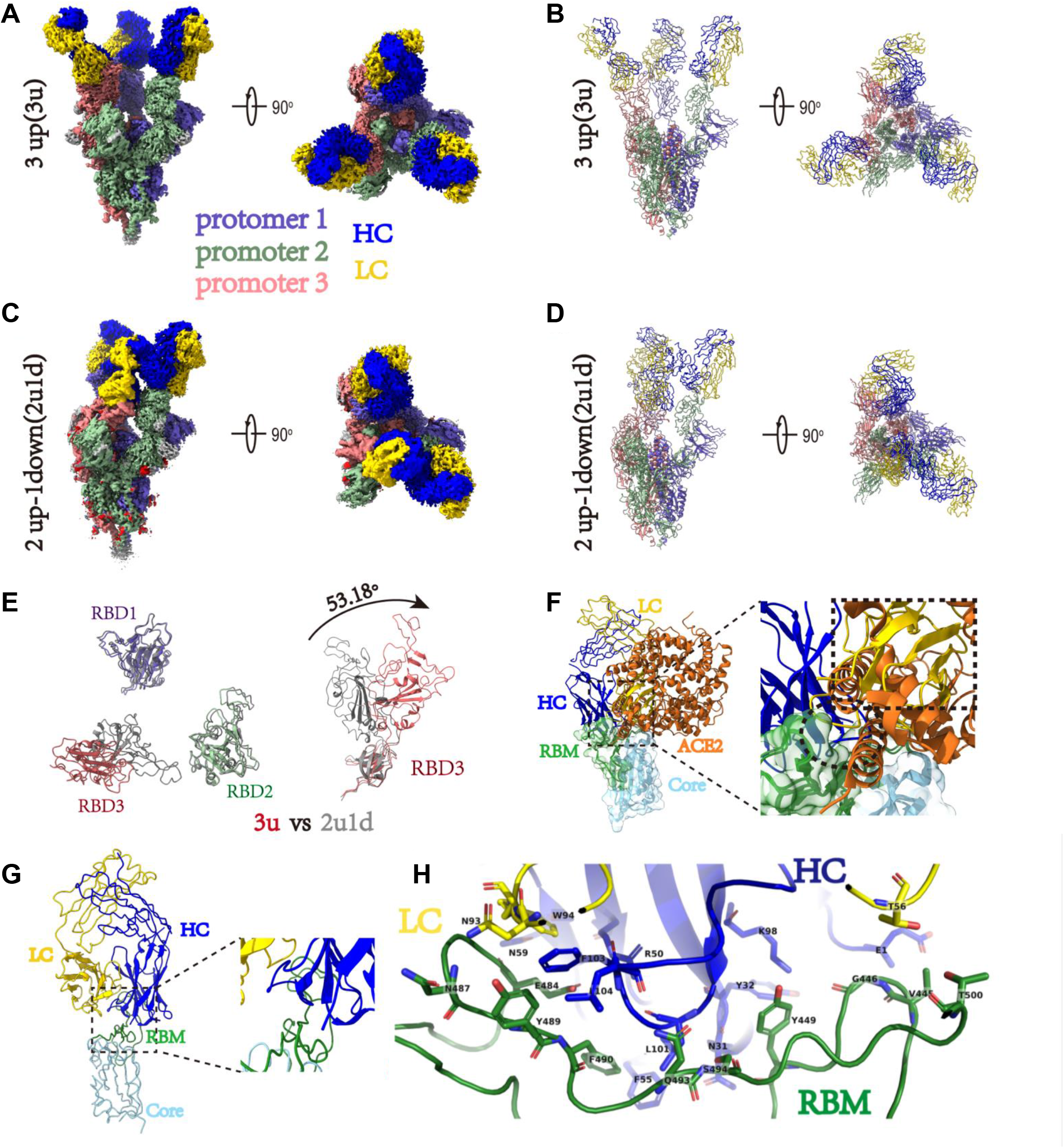
Cryo-EM structures of the SARS-CoV-2 S trimer in complex with B8-Fabs. **(A)** Side and top views of the Spike-B8 3u cryo-EM map showing 3 up RBDs each bound with a B8 Fab. Protomer 1, 2, and 3 are shown in slate blue, dark sea green, and India red, respectively. Heavy chain and light chain of the B8-Fab are in blue and gold, respectively. This color scheme was used throughout panels **(A)-(E)**. **(B)** Side and top views of the Spike-B8 3u cryo atomic model. **(C)** Side and top views of the Spike-B8 2u1d cryo-EM map showing two up RBDs up (RBD-1 and RBD-2) and one RBD down (RBD-3), each bound to a B8-Fab. **(D)** Side and top views of the Spike-B8 2u1d cryo atomic model. **(E)** Structural comparation of RBDs between Spike-B8 3u (different colors) and Spike-B8 2u1d (gray). **(F)** ACE2 (chocolate color, PDB: 6M0J) may clash with the heavy chain (blue) and light chain (gold) of the B8-Fab. ACE2 and the Fab share overlapping epitopes on the RBM (dotted black circle), and the framework of the B8-VL appears to clash with ACE2 (dotted black frame). The RBD core and RBM are shown in light sky blue and green, respectively. **(G)** Atomic model of an RBD-B8 complex portion in cartoon mode, shown with the same color scheme as in **(F)**. **(H)** The residues involved in interactions between B8 and the RBM. The heavy and light chain of the B8-Fab are in blue and gold, respectively. The RBM is shown in green.

## DISCUSSION

In this study, we investigated the preventive potential of a potent RBD-specific NAb B8 primarily in the forms of monomeric and dimeric IgA against live intranasal SARS-CoV-2 infection in the golden Syrian hamster model as compared with B8-IgG1 ^47,48^. While these B8-IgA antibodies maintained neutralizing activities against SARS-CoV-2 *in vitro* similar to those of B8-IgG1, they displayed distinct *in vivo* effects, with clear differences in their capacity to modulate viral infection in the NT. Pretreatment by intranasal administration of 4.5 mg/kg of monomeric B8-mIgA1 or B8-mIgA2 did not significantly reduce infectious virus production in the NT homogenates. On the contrary, the antibody isotype had a marked effect, as intranasal administration of 4.5 mg/kg and 13.5 mg/kg dimeric B8-dIgA1 or B8-dIgA2 paradoxically increased the amount of infectious virus (PFU) in NT homogenates. This enhancing effect was not observed with several intranasal IgG HuNAbs previously tested by other groups or our team ^47,54^. Mechanistically, instead of neutralization, virus-bound B8-dIgA1 and B8-dIgA2 used CD209 as an alternative receptor to infect non-ACE2 cells. CD209^+^ cells were increased and permissive to viral infection in the olfactory epithelium of Syrian hamsters upon SARS-CoV-2 infection, suggesting that this cell population could contribute to viral mucosal seeding. Indeed, we found that CD209 expressing cells could be infected *in vitro* by live SARS-CoV-2 at 0.05 MOI in the presence of B8-dIgA1 and B8-dIgA2. Since none of the B8-based MAbs could prevent SARS-CoV-2 cell-to-cell transmission, even at high concentration *in vitro*, virus-laden mucosal CD209^+^ cells might trans-infect ACE2^+^ cells through cell-to-cell contacts in NT, resulting in enhanced infection and injury. Cryo-EM analysis further indicated that B8 is a typical class II HuNAb that binds to the SRAS-CoV-2 spike RBD in either a 3u or a 2u1d mode. Our findings, therefore, reveal a previously unrecognized pathway for RBD-specific dimeric IgA-mediated enhancement of SARS-CoV-2 nasal infection and injury in Syrian hamsters.

The role of dimeric IgA has been explored primarily for mucosal transmitted viruses. At the mucosal surface, the major IgA type is the secretory form, which is generated from dIgA by the acquisition of a secretory component upon endocytosis and secretion by epithelial cells. In the simian AIDS macaque model, neutralizing dIgA given directly into the rectal lumen can prevent viral acquisition in rhesus monkeys challenged via the mucosal route ^55^. Although the administered dIgA did not contain the secretory component (SC), they might have associated with free SC, which is present in mucosal secretions such as human lung lavages ^56^. Neutralizing dIgA1 and dIgA2 could be protective through several mechanisms, including direct virus neutralization, virion capture, or the inhibition of virion transcytosis across the epithelium ^40^. In this macaque study, however, Watkins et al. demonstrated that the dimeric HGN194 dIgA2 protected only 1/6 animals in a rectal challenge model ^55^. Recently, Taylor et al. found an increase in virion number and penetration depth in the transverse colon and mesenteric lymph nodes, after mucosal treatment with the HGN194 dIgA2 compared to a PBS control ^57^. The authors suggested that virus-specific dIgA somehow mediated the delivery of virus immune complexes to the mesenteric lymph nodes for systemic infection. Here, we report that SARS-CoV-2 may subvert the action of potent neutralizing antibodies, as pretreatment with neutralizing B8-dIgA1 and B8-dIgA2 induced a more robust nasal infection via a previously unrecognized mode of viral enhancement.

SARS-CoV-2 engages CD209^+^ cells to evade ACE2-dependent neutralizing B8-dIgA1 and B8-dIgA2 for enhanced NT infection and injury. Previous studies have indicated various scenarios for ADE occurrence in viral infections. The well-known dengue ADE has been associated with poorly neutralizing cross-reactive antibodies against a heterologous viral serotype, leading to increased infection of FcγR-expressing cells ^58^. Recent findings suggested that an increase in afucosylated antibodies contribute to dengue ADE ^59^. In contrast, vaccine-associated enhanced respiratory disease induced by respiratory syncytial virus has not been found to be antibody-dependent ^60^. For SARS and MERS, ADE observed *in vitro* depended on binding of the antibody Fab to the virus and the binding of the Fc component to FcγR on target cells ^61^. One study found that spike IgG antibody abrogated wound-healing responses in SARS-CoV-1-infected Chinese macaques ^62^. In the case of COVID-19, vaccination and passive immunization studies have not revealed ADE of disease severity ^63^. Comprehensive studies, however, are necessary to define the clinical correlates of protective immunity against SARS-CoV-2, especially in the context of vaccine breakthrough infections. During natural infection, one study indicated that the increase in afucosylated antibodies might contribute to COVID-19 severity ^64^. To date, four classes of potent HuNAbs have been isolated from convalescent COVID-19 patients ^34,53^. The molecular mechanism of neutralization for most potent HuNAbs was primarily through blocking the interaction between ACE2 and the spike RBD. Currently, systemic RBD-specific HuNAb treatment remains to be improved for therapeutic suppression of SARS-CoV-2 replication in the NT or URT, both in animal models and human trials ^47,48,65^. One limitation is the insufficient amounts of HuNAbs distributed on the nasal mucosal surface for protection ^47^. Other reasons might include alternative entry pathways engaged by SARS-CoV-2 to evade HuNAbs. To this end, Liu et al. reported recently that antibodies against the spike N-terminal domain (NTD) induced an open conformation of the RBD and thus enhanced the binding capacity of the spike to the ACE2 receptor, leading to increased viral infectivity ^66^. Yeung et al. demonstrated nicely that SARS-CoV-2 could engage soluble ACE2 (sACE2) and then bind alternate receptors for viral entry, through interaction between a spike/sACE2 complex with the angiotensin II AT1 receptor, or interaction between a spike/sACE2/vasopressin complex with the AVPR1B vasopressin receptor, respectively ^49^. In this study, we found that, in the presence of potent neutralizing B8-dIgA1 or B8-dIgA2 antibodies, SARS-CoV-2 used the cellular receptor CD209 for capture or infection, which likely expanded the use of CD209^+^ cells as target cells, leading to enhanced NT infection and trans-infection. Interestingly, a preprint report suggests that cells expressing CD209 can be infected directly by SARS-CoV-2 through an interaction of the spike with the NTD instead of the RBD ^67^. This mode of action, however, was unlikely to explain our findings, because no enhancement of SARS-CoV-2 nasal infection was found in presence of control dIgA1 and dIgA2. Our results rather suggest that the direct binding of virus-bound B8-dIgA1 or virus-bound B8-dIgA2 to CD209 is a likely pathway, resulting in the more severe SARS-CoV-2 nasal infection and damage. In line with our results, a previous study demonstrated that dIgA itself can use CD209 as a cellular receptor ^52^. During the entry process, since neither B8-dIgA1 nor B8-dIgA2 could prevent virus cell-to-cell transmission, infected mucosal CD209^+^ cells might enable a more robust viral transmission to ciliated nasal epithelial cells in NT, which show the highest expression of ACE2 and TMPRSS2 receptors ^65^. In support of this notion, previous studies indicated that mucosal DCs can capture HIV-1 through binding of its envelope glycoproteins to CD209 and efficiently transfer the bound virions to CD4^+^ T cells, in a process called trans-enhancement or trans-infection ^68^. The trans-infection markedly decreased the neutralization efficiency of potent NAbs directed at HIV ^69^. Moreover, although monocyte-derived DCs (MDDCs) cannot support productive SARS-CoV-2 replication ^70^, a recent study demonstrated that MDDCs could mediate efficient viral trans-infection of the Calu-3 human respiratory cell line ^71^. Our findings of increased number of infectious viruses in NT, therefore, have significant implications for SARS-CoV-2 transmission, COVID-19 pathogenesis, and immune interventions.

### Limitations of the study

As our *in vivo* findings were obtained in Syrian hamsters, it remains to be determined whether CD209^+^ DCs are abundantly recruited to the nasal mucosa in SARS-CoV-2-infected humans. It is, however, known that myeloid DCs are increased in the nasal epithelium upon infection ^51,72^. Our preliminary analysis of the human nasal cytology data (under accession code EGAS00001004082) revealed the presence of increased CD209^+^ DCs in addition to abundant ACE2, TMPRSS2, and furin expression in the apical side of multiciliated cells of SARS-CoV-2-infected human subjects (Supplementary Fig. 9) ^73^. Another limitation is that we did not have an NTD-specific neutralizing dIgA for comparison. Besides the class II neutralizing dIgA such as B8-mIgA2 and B8-dIgA2 used in this study, neutralizing dIgA belonging to other classes should also be investigated in future. We also do not know whether other cellular receptor such as the polymeric immunoglobulin receptor (pIgR) plays a role in B8-dIgA-enhanced SARS-CoV-2 nasal infection. The pIgR is responsible for transcytosis of soluble dIgAs and immune complexes from the basolateral to the apical epithelial surface. It remains uncertain whether B8-dIgA-enhanced NT infection would lead to worse neuro-COVID-19. Because current intramuscular vaccinations might not induce secretory dIgA at the nasal mucosal sites ^74^, it remains unknown whether the dIgA-mediated ADE would happen in people who received the COVID-19 vaccines or dIgA treatment. Future studies are needed to address these limitations.

## METHODS

### Human subjects

A total of 4 patients with COVID-19 including 3 convalescent cases and one acute case were recruited between February and May 2020. All patients were confirmed by reverse-transcription polymerase chain reaction (RT-PCR) as described previously ^28^. Clinical and laboratory findings were entered into a predesigned database. Written informed consent was obtained from all patients. This study was approved by the Institutional Review Board of The University of Hong Kong/Hospital Authority Hong Kong West Cluster, the Hong Kong East Cluster Research Ethics Committee, and the Kowloon West Cluster Research Ethics Committee (UW 13-265, HKECREC-2018-068, KW/EX-20-038[144-26]).

### Syrian hamsters

The animal experimental plan was approved by the Committee on the Use of Live Animals in Teaching and Research (CULATR 5359-20) of the University of Hong Kong (HKU). Male and female golden Syrian hamsters (*Mesocricetus auratus*) (aged 6–10 weeks) were purchased from the Chinese University of Hong Kong Laboratory Animal Service Centre through the HKU Laboratory Animal Unit (LAU). The animals were kept in Biosafety Level-2 housing and given access to standard pellet feed and water ad libitum following LAU’s standard operational procedures (SOPs). The viral challenge experiments were then conducted in our Biosafety Level-3 animal facility following SOPs strictly, with strict adherence to SOPs

### Cell lines

HEK293T cells, HEK293T-hACE2 cells Vero-E6 cells, HK2 cells and Vero-E6-TMPRSS2 cells were maintained in DMEM containing 10% FBS, 2 mM L-glutamine, 100 U/mL/mL penicillin and incubated at 37 □ in a 5% CO_2_ setting ^62^. Expi293F™ cells were cultured in Expi293TM Expression Medium (Thermo Fisher Scientific) at 37 □ in an incubator with 80% relative humidity and a 5% CO_2_ setting on an orbital shaker platform at 125 ±5 rpm/min (New Brunswick innova™ 2100) according to the manufacturer’s instructions.

### ELISA analysis of plasma and antibody binding to RBD and trimeric spike

The recombinant RBD and trimeric spike proteins derived from SARS-CoV-2 (Sino Biological) were diluted to final concentrations of 1 μg/mL/mL, then coated onto 96-well plates (Corning 3690) and incubated at 4 °C overnight. Plates were washed with PBS-T (PBS containing 0.05% Tween-20) and blocked with blocking buffer (PBS containing 5% skim milk or 1% BSA) at 37 °C for 1 h. Serially diluted plasma samples or isolated monoclonal antibodies were added to the plates and incubated at 37 °C for 1 h. Wells were then incubated with a secondary goat anti-human IgG labelled with horseradish peroxidase (HRP) (Invitrogen) or with a rabbit polyclonal anti-human IgA alpha-chain labelled with HRP (Abcam) and TMB substrate (SIGMA). Optical density (OD) at 450 nm was measured by a spectrophotometer. Serially diluted plasma from healthy individuals or previously published monoclonal antibodies against HIV-1 (VRC01) were used as negative controls.

### Isolation of RBD-specific IgG+ single memory B cells by FACS

RBD-specific single B cells were sorted as previously described ^75^. In brief, PBMCs from infected individuals were collected and incubated with an antibody cocktail and a His-tagged RBD protein for identification of RBD-specific B cells. The cocktail consisted of the Zombie viability dye (Biolegend), CD19-Percp-Cy5.5, CD3-Pacific Blue, CD14-Pacific Blue, CD56-Pacific Blue, IgM-Pacific Blue, IgD-Pacific Blue, IgG-PE, CD27-PE-Cy7 (BD Biosciences) and the recombinant RBD-His described above. Two consecutive staining steps were conducted: the first one used an antibody and RBD cocktail incubation of 30 min at 4 °C; the second staining involved staining with anti-His-APC and anti-His-FITC antibodies (Abcam) at 4 °C for 30 min to detect the His tag of the RBD. The stained cells were washed and resuspended in PBS containing 2% FBS before being strained through a 70-μm cell mesh filter (BD Biosciences). RBD-specific single B cells were gated as CD19^+^CD27^+^CD3^−^CD14^−^ CD56^−^IgM^−^IgD^−^IgG^+^RBD^+^ and sorted into 96-well PCR plates containing 10 μL of RNAase-inhibiting RT-PCR catch buffer (1M Tris-HCl pH 8.0, RNase inhibitor, DEPC-treated water). Plates were then snap-frozen on dry ice and stored at −80 °C until the reverse transcription reaction.

### Single B cell RT-PCR and antibody cloning

Single memory B cells isolated from PBMCs of infected patients were cloned as previously described ^76^. Briefly, one-step RT-PCR was performed on sorted single memory B cell with a gene specific primer mix, followed by nested PCR amplifications and sequencing using the heavy chain and light chain specific primers. Cloning PCR was then performed using heavy chain and light chain specific primers containing specific restriction enzyme cutting sites (heavy chain, 5′-AgeI/3′-SalI; kappa chain, 5′-AgeI/3′-BsiWI). The PCR products were purified and cloned into the backbone of antibody expression vectors containing the constant regions of human Igγ1 or Igα1 and Igα2. The Igα1 and Igα2 vectors were purchased from InvivoGen (pfusess-hcha1 for IgA1 and pfusess-hcha2m1 for IgA2). The constructed plasmids containing paired heavy and light chain expression cassettes were co-transfected into 293T cells (ATCC) grown in 6-well plates. Antigen-specific ELISA and pseudovirus-based neutralization assays were used to analyze the binding capacity to SARS-CoV-2 RBD and the neutralization capacity of transfected culture supernatants, respectively.

### Genetic analysis of the BCR repertoire

Heavy chain and light chain germline assignment, framework region annotation, determination of somatic hypermutation (SHM) levels (in nucleotides) and CDR loop lengths (in amino acids) were performed with the aid of the IMGT/HighV-QUEST software tool suite (https://www.imgt.org/HighV-QUEST). Sequences were aligned using Clustal W in the BioEdit sequence analysis package (Version 7.2). Antibody clonotypes were defined as a set of sequences that share genetic V and J regions as well as an identical CDR3.

### Antibody production and purification

The paired antibody VH/VL chains were cloned into Igγ, Igα1 or Igα2 and Igk expression vectors using T4 ligase (NEB). For production of IgG and monomeric IgA, the plasmids with paired heavy chain (IgG, IgA1, IgA2) and light chain genes were co-transfected into Expi293™ expression system (Thermo Fisher Scientific) following the manufacturer’s protocol to produce recombinant monoclonal antibodies. For dIgA antibody production, plasmids of paired heavy chain (IgA1, IgA2) and kappa light chain together with a J chain were co-transfected into Expi293™ expression system (Thermo Fisher Scientific) at the ratio of 1:1:1 following the manufacturer’s instructions. Antibodies produced from cell culture supernatants were purified immediately by affinity chromatography using recombinant Protein G-Agarose (Thermo Fisher Scientific) or CaptureSelect™ IgA Affinity Matrix (Thermo Fisher Scientific) according to the manufacturer’s instructions, to purify IgG and IgA, respectively. The purified antibodies were concentrated by an Amicon ultracentrifuge filter device (molecular weight cut-off 10 kDa; Millipore) to a volume of 0.2 mL in PBS (Life Technologies), and then stored at 4 °C or −80 °C for further characterization.

### Size exclusion chromatography

The prepacked HiLoad 26/60 Superdex™ 200pg (code No. 17-1071-01, Cytiva) column was installed onto the Amersham Biosciences AKTA FPLC system. After column equilibration with 2 column volumes (CV) of PBS, the concentrated IgA antibodies were applied onto the column using a 500-ul loop at a flow rate of 2 mL/min. Dimers of IgA1 or IgA2 were separated from monomers upon washing with 2 CV of PBS. The milli-absorbance unit at OD280nm was recorded during the washing process. 2 mL-fractions were collected, pooled, concentrated and evaluated by western blot using mouse anti-IGJ monoclonal antibody [KT109] (Abcam) and rabbit anti-human IgA alpha chain antibody (Abcam).

### Pseudovirus-based neutralization assay

The neutralizing activity of NAbs was determined using a pseudotype-based neutralization assay as we previously described ^77^. Briefly, The pseudovirus was generated by co-transfection of 293T cells with pVax-1-S-COVID19 and pNL4-3Luc_Env_Vpr, carrying the optimized spike (S) gene (QHR63250) and a human immunodeficiency virus type 1 backbone, respectively ^77^. Viral supernatant was collected at 48 h post-transfection and frozen at −80 °C until use. The serially diluted monoclonal antibodies or sera were incubated with 200 TCID_50_ of pseudovirus at 37 °C for 1 hour. The antibody-virus mixtures were subsequently added to pre-seeded HEK 293T-ACE2 cells. 48 hours later, infected cells were lysed to measure luciferase activity using a commercial kit (Promega, Madison, WI). Half-maximal (IC_50_) or 90% (IC_90_) inhibitory concentrations of the evaluated antibody were determined by inhibitor vs. normalized response -- 4 Variable slope using GraphPad Prism 6 or later (GraphPad Software Inc.).

### Neutralization activity of monoclonal antibodies against authentic SARS-CoV-2

The SARS-CoV-2 focus reduction neutralization test (FRNT) was performed in a certified Biosafety level 3 laboratory. Neutralization assays against live SARS-CoV-2 were conducted using a clinical isolate (HKU-001a strain, GenBank accession no: MT230904.1) previously obtained from a nasopharyngeal swab from an infected patient ^78^. The tested antibodies were serially diluted, mixed with 50 μL of SARS-CoV-2 (1×10^3^ focus forming unit/mL, FFU/mL) in 96-well plates, and incubated for 1 hour at 37°C. Mixtures were then transferred to 96-well plates pre-seeded with 1×10^4^/well Vero E6 cells and incubated at 37°C for 24 hours. The culture medium was then removed and the plates were air-dried in a biosafety cabinet (BSC) for 20 mins. Cells were then fixed with a 4% paraformaldehyde solution for 30 min and air-dried in the BSC again. Cells were further permeabilized with 0.2% Triton X-100 and incubated with cross-reactive rabbit sera anti-SARS-CoV-2-N for 1 hour at RT before adding an Alexa Fluor 488 goat anti-rabbit IgG (H+L) cross-adsorbed secondary antibody (Life Technologies). The fluorescence density of SARS-CoV-2 infected cells were scanned using a Sapphire Biomolecular Imager (Azure Biosystems) and the neutralization effects were then quantified using Fiji software (NIH).

### Antibody binding kinetics, and competition with the ACE2 receptor measured by Surface Plamon Resonance (SPR)

The binding kinetics and affinity of recombinant monoclonal antibodies for the SARS-CoV-2 spike protein (ACROBiosystems) were analysed by SPR (Biacore 8K, GE Healthcare). Specifically, the spike protein was covalently immobilized to a CM5 sensor chip via amine groups in 10mM sodium acetate buffer (pH 5.0) for a final RU around 500. SPR assays were run at a flow rate of 30 mL/min in HEPES buffer. For conventional kinetic/dose-response, serial dilutions of monoclonal antibodies were injected across the spike protein surface for 180s, followed by a 600s dissociation phase using a multi-cycle method. Remaining analytes were removed in the surface regeneration step with the injection of 10 mM glycine-HCl (pH 2.0) for 2×30s at a flow rate of 30 μl/min. Kinetic analysis of each reference subtracted injection series was performed using the Biacore Insight Evaluation Software (GE Healthcare). All sensorgram series were fit to a 1:1 (Langmuir) binding model of interaction. Before evaluating the competition between antibodies and the human ACE2 peptidase domain, both the saturating binding concentrations of antibodies and of the ACE2 protein (ACROBiosystems) for the immobilized SARS-CoV-2 spike protein were determined separately. In the competitive assay, antibodies at the saturating concentration were injected onto the chip with immobilized spike protein for 120s until binding steady-state was reached. ACE2 protein also used at the saturating concentration was then injected for 120s, followed by another 120s of injection of antibody to ensure a saturation of the binding reaction against the immobilized spike protein. The differences in response units between ACE2 injection alone and prior antibody incubation reflect the antibodies’ competitive ability against ACE2 binding to the spike protein.

### Hamster experiments

*In vivo* evaluation of monoclonal antibody B8-IgG1, B8-mIgA1, B8-mIgA2, B8-dIgA1, B8-dIgA2 in the established golden Syrian hamster model of SARS-CoV-2 infection was performed as described previously, with slight modifications ^48^. Approval was obtained from the University of Hong Kong (HKU) Committee on the Use of Live Animals in Teaching and Research. Briefly, 6-8-week-old male and female hamsters were housing with access to standard pellet feed and water *ad libitum* until live virus challenge in the BSL-3 animal facility. The hamsters were randomized from different litters into experimental groups. Experiments were performed in compliance with the relevant ethical regulations ^48^. For prophylaxis studies, 24 hours before live virus challenge, three groups of hamsters were intraperitoneally or intranasally administered with one dose of test antibody in phosphate-buffered saline (PBS) at the indicated dose. At day 0, each hamster was intranasally inoculated with a challenge dose of 100 μL of Dulbecco’s Modified Eagle Medium containing 10^5^ PFU of SARS-CoV-2 (HKU-001a strain, GenBank accession no: MT230904.1) under anaesthesia with intraperitoneal ketamine (200 mg/kg) and xylazine (10 mg/kg). For pre-treatment study, each hamster received one 1.5 mg/kg dose of intraperitoneal B8-IgG1 at 24, 48, 72 hours (n=4 per group) after virus challenge. The hamsters were monitored twice daily for clinical signs of disease. Syrian hamsters typically clear virus within one week after SARS-CoV-2 infection. Accordingly, animals were sacrificed for analysis at day 4 after virus challenge with high viral loads ^48^. Half the nasal turbinate, trachea, and lung tissues were used for viral load determination by quantitative SARS-CoV-2-specific RdRp/Hel RT-qPCR assay ^28^ and infectious virus titration by plaque assay ^48^.

### Cryo-EM sample preparation and data acquisition

The purified SARS-CoV-2 S-B8 protein complexes were concentrated before being applied to the grids. Aliquots (4 μL) of the protein complex were placed on glow-discharged holey carbon grids (Quantifoil Au R1.2/1.3, 300 mesh). The grids were blotted and flash-frozen in liquid ethane cooled by liquid nitrogen with a Vitrobot apparatus (Mark IV, ThermoFisher Scientific). The grids sample quality was verified with an FEI Talos Arctica 200-kV electron microscope (Thermo Fisher Scientific). The verified grids with optimal ice thickness and particle density were transferred to a Titan Krios operating at 300 kV and equipped with a Cs corrector, a Gatan K3 Summit detector (Gatan Inc.) and a GIF Quantum energy filter (slit width 20 eV). Micrographs were recorded in the super-resolution mode with a calibrated pixel size of 0.54895 Å. Each movie has a total accumulated exposure of 50 e−/A□2 fractionated in 32 frames. The final image was binned 2-fold to a pixel size of 1.0979 A□. AutoEMation was used for the fully automated data collection. The defocus value of each image, which was set from −1.0 to −2.0 μm during data collection, was determined by Gctf. Data collection statistics are summarized in Supplementary Table 11.

### Cryo-EM data processing

The procedure for image processing of SARS-CoV-2 S-B8 complex is presented in Supplementary Fig. 2. In brief, Motion Correction (MotionCo2), CTF-estimation (GCTF) and non-templated particle picking (Gautomatch, http://www.mrc-lmb.cam.ac.uk/kzhang/) were automatically executed by the TsinghuaTitan.py program (developed by Dr. fang Yang). Sequential data processing was carried out on RELION 3.0 and RELION 3.1. Initially, 2,436,776 particles were auto-picked by Gautomatch or RELION 3.0 from 4213 micrographs. After several 2D classifications, 1,451,176 particles were selected and applied for 3D classification with one class. Two different states were obtained after further 3D classification: 3 RBD in up conformation bound with B8 Fab (3u), and 2 up RBDs and 1 down RBD with each bound to a B8 Fab (2u1d). 616,799 particles for the 2u1d state and 351,095 particles for the 3u state were subjected to 3D auto-refinement, yielding final resolutions at 3.21 A□ and 3.06 A□, respectively. Further CTF refinement and Bayesian polishing improved the resolution to 2.65 Å (2u1d, C1 symmetry) and 2.67 Å (3u, C3 symmetry) with better map quality. To improve the RBD-B8 portion map density, focused local search classification was applied for each RBD-B8 portion with an adapted soft mask. The best classes for each RBD-Fab portion were selected and yielded a final resolution at 3.56 Å (RBD-Fab1, up), 3.34 Å (RBD-Fab2, up), 3.69 Å (RBD-Fab3, down), 3.87 Å (RBD-Fab3, up) from 479,305, 508,653, 656,429, and 136,482 particles, respectively. Further CTF refinement and Bayesian polishing improved the resolution of RBD-Fab2 to 3.11 Å with better map quality. RBD-Fab maps were fitted onto the whole structure map using Chimera, then combined using PHENIX combine_focused_maps. The reported resolutions were estimated with the gold-standard Fourier shell correlation (FSC) cutoff of 0.143 criterion. Data processing statistics are summarized in Supplementary Table 11.

### Model building and structure refinement

The spike model (PDB code: 6VSB) and the initial model of the B8 Fab generated by SWISS-Model were fitted into the EM density map, and further manually adjusted with Coot. Glocusides were built manually with carbohydrate tool in Coot. The atomic models were refinement using Phenix in real space with secondary structure and geometry restraints. The final structures were validated using Phenix.molprobity. UCSF Chimera, ChimeraX and PyMol were used for map segmentation and figure generation. Model refinement statistics are summarized in Supplementary Table 11.

### SARS-CoV-2 infection of reconstructed human nasal epithelia

MucilAir™, corresponding to reconstructed human nasal epithelium cultures differentiated *in vitro* for at least 4 weeks, were purchased from Epithelix (Saint-Julien-en-Genevois, France). The cultures were generated from pooled nasal tissues obtained from 14 human adult donors. Cultures were maintained in air/liquid interface (ALI) conditions in transwells with 700 μL of MucilAir™ medium (Epithelix) in the basal compartment, and then kept at 37 °C under a 5% CO2 atmosphere. SARS-CoV-2 infection was performed as previously described ^50^. Briefly, the apical side of ALI cultures was washed 20 min at 37 °C in Mucilair™ medium to remove mucus. Cells were then incubated with 10^4^ plaque-forming units (PFU) of the isolate BetaCoV/France/IDF00372/2020 (EVAg collection, Ref-SKU: 014V-03890; kindly provided by S. Van der Werf). The viral input was diluted in DMEM medium to a final volume 100 μL, and then left on the apical side for 4 h at 37 °C. Control wells were mock treated with DMEM medium (Gibco) for the same duration. Viral inputs were removed by washing twice with 200 μL of PBS (5 min at 37 °C) and once with 200 μL Mucilair™ medium (20 min at 37 °C). The basal medium was replaced every 2-3 days. Apical supernatants were harvested every 2-3 days by adding 200 μL of Mucilair™ medium on the apical side, with an incubation of 20 min at 37 °C prior to collection. For IgA treatment, cultures were washed once and then pretreated with antibodies added to the apical compartment for 1 h in 50μL. Viral input was then directly added to reach a final volume of 100 μL. The antibodies were added again at day 2 d.p.i. in the apical compartment during an apical wash (20 min at 37 °C). To test the effect of dIgA treatment in the presence of mucus, dIgA were added directly to the apical compartment of MucilAir™ cultures without an initial wash. After IgA treatment for 1h, the virus was added directly to the IgA/mucus mixture and left on the apical side for 4h at 37°C. After viral inoculation, a single brief wash was made to remove the viral input while limiting mucus loss. The cultures were then maintained as in the no-mucus condition.

### Viral RNA quantification in reconstructed human nasal epithelia

Apical supernatants were collected, stored at −80 °C until thawing and were then diluted 4-fold in PBS in a 96-well plate. Diluted supernatants were inactivated for 20 min at 80 °C. For SARS-CoV-2 RNA quantification, 1 μL of diluted supernatant was added to 4 μL of PCR reaction mix. PCR was carried out in 384-well plates using the Luna Universal Probe One-Step RT-qPCR Kit (New England Biolabs) with SARS-CoV-2 NP-specific primers (Forward 5’-TAA TCA GAC AAG GAA CTG ATT A-3’; Reverse 5’-CGA AGG TGT GAC TTC CATG-3’) on a QuantStudio 6 Flex thermocycler (Applied Biosystems). A standard curve was established in parallel using purified SARS-CoV-2 viral RNA.

### Histopathology and immunofluorescence (IF) staining

The lung and nasal turbinate tissues collected at necropsy were fixed in zinc formalin and then processed into paraffin-embedded tissue blocks. The tissue sections (4 μm) were stained with hematoxylin and eosin (H&E) for light microscopy examination as previously described with modifications ^47^. For identification and localization of SARS-CoV-2 nucleocapsid protein (NP) in organ tissues, immunofluorescence staining was performed on deparaffinized and rehydrated tissue sections using a rabbit anti-SARS-CoV-2-NP protein antibody together with an AF488-conjugated anti-rabbit IgG (Jackson ImmunoResearch, PA, USA). Briefly, the tissue sections were first treated with antigen unmasking solution (Vector Laboratories) in a pressure cooker. After blocking with 0.1% Sudan black B for 15 min and 1% bovine serum albumin (BSA)/PBS at RT for 30 min, the primary rabbit anti-SARS-CoV-2-NP antibody (1:4000 dilution with 1% BSA/PBS) was incubated at 4°C overnight. This step was followed by incubation with a FITC-conjugated donkey anti-rabbit IgG (Jackson ImmunoResearch) for 30 min and the sections were then mounted in medium with 4’,6-diamidino-2-phenylindole (DAPI). For identification of DC-SIGN expression, we stained the NT slices with rabbit anti-DC-SIGN primary antibody (Abcam) and Alexa Fluor 488 goat anti-rabbit IgG (H+L) cross-adsorbed secondary antibody (Life Technologies) according to the manufacturer’s instructions. For identification of ACE2 expression, the goat anti-ACE2 primary antibody (R&D) and Alexa Fluor 568 donkey anti-goat IgG (H+L) secondary antibodies (Invitrogen) according to the manufacturer’s instructions. All tissue sections were examined, and the fluorescence images and whole section scanning were captured using 5×, 10× and 20× objectives with Carl Zeiss LSM 980. NP^+^ cells per field were quantified based on the mean fluorescence intensity (MFI) using the ZEN BLACK 3.0 and ImageJ (NIH).

### Effects of B8-dIgA on SARS-CoV-2 infection in HK2 cells

HK2 cells were seeded into 24-well plates at the 40-50% confluency and cultured overnight. The B8-dIgA or control dIgA at the concentration of 1, 10, 100, 1000 ng/ml/mL and then mixed with SARS-CoV-2 (1:10 TCID50) and incubated for 1 hour at room temperature. The antibody/virus mixture was then added to HK2 cells after the cell culture medium was removed and washed with PBS once and incubated for 1 hour at 37°C. The infectious medium was replaced with fresh medium containing respective concentration of antibody after washing 3 times with PBS. 24 h later, the infected cells were imaged under fluorescence microscope after staining with AF488-conjugated anti-SARS-CoV-2 NP antibody. Alternatively, the infected cells were lysed and blotted for SARS-CoV-2 NP protein to determine the extent of infection. Tubulin was blotted as the internal control.

### B8-dIgA mediated enhancement via CD209

HEK293T cells were seeded into 10-cm dish at 40% confluency and cultured overnight. The HEK293T cells were transfected with human CD209 (Sino Biological) at 70%-90% confluency. The expression of CD209 was measured by flow cytometry. The transfected HEK293T-CD209 cells were seeded into 96-well plates with 2.4×10^4^ cells per well and cultured overnight. The HEK293T-CD209 cells were pre-treated with 10 ng/ml/mL of B8-dIgA or control dIgA and incubated for 6 h prior SARS-CoV-2 infection (MOI=0.05). 24 h later, cells were then fixed with 4% paraformaldehyde solution for 30 min and air-dried in the BSC. Cells were further permeabilized with 0.2% Triton X-100 and incubated with cross-reactive rabbit sera anti-SARS-CoV-2-N for 1 hour at RT before adding Alexa Fluor 488 goat anti-rabbit IgG (H+L) cross-adsorbed secondary antibody (Life Technologies). The fluorescence density of SARS-CoV-2 infected cells was acquired using a Sapphire Biomolecular Imager (Azure Biosystems) and then the MFI of four randomly selected areas of each sample was quantified using Fiji software (NIH).

### Effects of B8 antibodies on SARS-CoV-2 mediated cell-cell fusion

Vero-E6 TMPRSS2 cells were seeded into 48-well plates and cultured overnight. After treatment with B8 antibodies at the dose of 3000 ng/ml/mL for 1 hour, HEK293T cells transfected with SARS-CoV-2 spike-GFP were added into the treated Vero-E6 TMPRSS2 cells and co-cultured for 48 hours. The cell-cell fusion between Vero-E6 TMPRSS2 and HEK293T-Spike-GFP was then determined under a fluorescence microscope (Nikon ELIPSE) and the images of randomly selected region were captured using 4× and 10× objectives using the Nikon software.

### Re-analysis of published nasal brushing single-cell data

The preprocessed scRNA-seq data from nasal brushing samples of 2 healthy controls and 4 COVID-19 patients were downloaded from Gene Expression Omnibus (GEO) database with accession numbers GSE171488 and GSE164547. Quality control metrics were consistent with the original article [PMID: 34003804] and performed based on the R package Seurat (version 4.0.3) [PMID: 34062119]. Harmony [PMID: 31740819] was used to integrate the samples based on the top 4000 most variable genes obtained with the FindVariableFeatures() function in Seurat. CD14^+^ (monocyte) cells were extracted for further analysis. The annotation of the cell type was performed by manually checking the marker genes of each cluster identified from the FindAllMarkers() function in Seurat.

### Quantification and statistical analysis

Statistical analysis was performed using PRISM 6.0 or later. Ordinary one-way ANOVA and multiple comparisons were used to compare group means and differences between multiple groups. Unpaired Student’s *t* tests were used to compare group means between two groups only. A P-value <0.05 was considered significant. The the number of independent experiments performed, the number of animals in each group, and the specific details of statistical tests are reported in the figure legends and the Methods section.

## Supporting information

Supplemental meterials

## SUPPLEMENTAL INFORMATION

The supplemental information includes 13 Tables and 9 Figures.

## ACKNOWLEDGMENTS

We acknowledge financial supports from the Hong Kong Research Grants Council Collaborative Research Fund (C7156-20G to Z.C.); the National Key Research and Development Project of China (2020YFC0860600) and the National Program on Key Research Project of China (2020YFA0707500 and 2020YFA0707504); the Health and Medical Research Fund (COVID1903010-7), the Food and Health Bureau, The Government of the Hong Kong Special Administrative Region; Innovation and Technology Fund (ITF), The Government of the Hong Kong Special Administrative Region; the University Development Fund and Li Ka Shing Faculty of Medicine Matching Fund from the University of Hong Kong to the AIDS Institute; the Consultancy Service for Enhancing Laboratory Surveillance of Emerging Infectious Diseases and Research Capability on Antimicrobial Resistance for Department of Health of the Hong Kong Special Administrative Region Government; Sanming Project of Medicine in Shenzhen, China (SZSM201911014); the High Level-Hospital Program, Health Commission of Guangdong Province, China; the research project of Hainan academician innovation platform (YSPTZX202004); and the Hainan talent development project (SRC200003); L.A.C’s team was supported by the Urgence COVID-19 Fundraising Campaign of Institute Pasteur (TROPICORO project). The study was also supported by generous donations of the Friends of Hope Education Fund, Lee Wan Keung Charity Foundation Limited, Shaw Foundation Hong Kong, Michael Seak-Kan Tong, Richard Yu and Carol Yu, May Tam Mak Mei Yin, Hong Kong Sanatorium & Hospital, Hui Ming, Hui Hoy and Chow Sin Lan Charity Fund Limited, Chan Yin Chuen Memorial Charitable Foundation, Marina Man-Wai Lee, the Hong Kong Hainan Commercial Association South China Microbiology Research Fund, the Jessie & George Ho Charitable Foundation, Perfect Shape Medical Limited, Kai Chong Tong, Tse Kam Ming Laurence, Foo Oi Foundation Limited, Betty Hing-Chu Lee, Ping Cham So, and Lo Ying Shek Chi Wai Foundation. Z.C.’s team was also partly supported by the Theme-Based Research Scheme (T11-706/18-N to Z.C.).

## AUTHOR CONTRIBUTIONS

Conceptualization, Z.C.; HuNAb cloning, Z.B.; experimental design, Z.C., Z.B., R.Z., J.F.W.C.; hamster experiments, J.F.W.C., C.C.S.C., V.K.M.P., C.C.Y.C., K.K.H.C., and JC; cryoEM study, J.Z., J.G., Z.Y.W., X.W.; SPR experiments, Q.Z., S.S. and L.Z.; LL confocal imaging, L.L. and D.Z.; MucilAir™ experiment, R.R. and L.A.C.; HK-2 experiment, M.L.Y.; nasal cytology data analysis, M.Y. and R.S.; clinical specimens, K.K.W.T.; *in vitro* experiments, H.C., Z.D., K.K.A., H.H., H.O.M., J.C., C.L., J.Z.; manuscript preparation, Z.C., Z.B., R.Z., L.Z., K.Y.Y.; study supervision, Z.C., K.Y.Y. and L.Z.

## DECLARATION OF INTEREST

J.F.W.C. has received travel grants from Pfizer Corporation Hong Kong and Astellas Pharma Hong Kong Corporation Limited and was an invited speaker for Gilead Sciences Hong Kong Limited and Luminex Corporation. The funding sources had no role in study design, data collection, analysis or interpretation or writing of the report. The other authors declare no conflicts of interest except for a provisional patent application filed for human monoclonal antibodies generated in our laboratory.

## Reporting Summary

Further information on research design is available in the Nature Research Reporting Summary linked to this article.

## Data availability

The data of this studies are available upon reasonable request and accession codes will be available before publication.

## Code availability

No custom computer code or algorithm used to generate results that are reported in the paper and central to its main claims.

