## Supplemental meterials for "SARS-CoV-2 hijacks neutralizing dimeric IgA for enhanced nasal infection and injury": Supplementary figures and tables 20210921 (Nature Micro).pdf

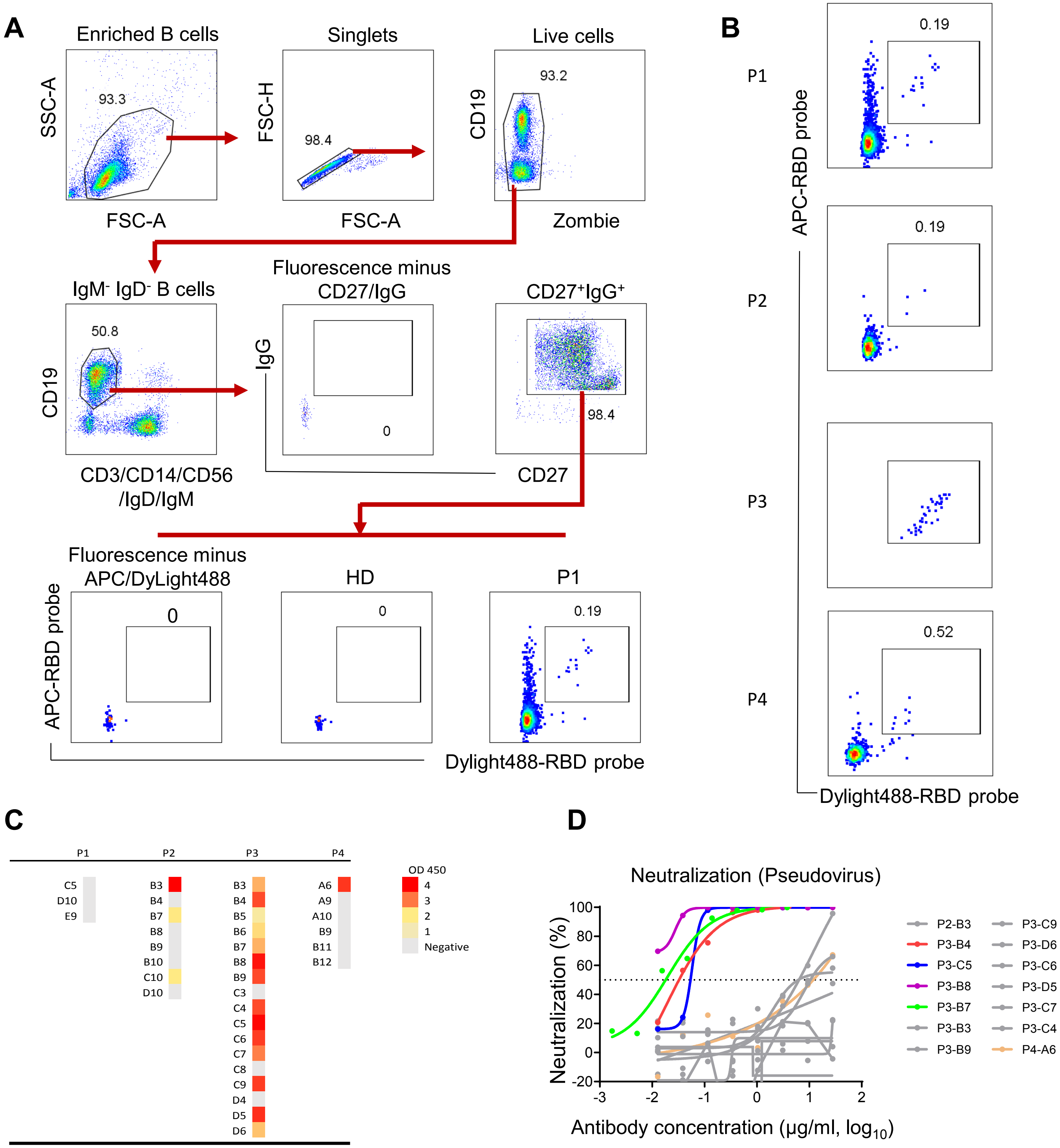

**Supplementary Fig. 1. Isolation of SARS-CoV-2 specific antibodies from sorted memory B cells.**

**(A)** The gating strategy for isolation of SARS-CoV-2 RBD-specific memory B cells by flow cytometry.

**(B)** The RBD double positive cell population was obtained from each subject.

**(C)** RBD-binding response of individual monoclonal antibodies from 4 subjects by ELISA. The colour scale indicated the absorbance value at OD450 nm.

**(D)** Neutralization activity was determined for screened antibodies against SARS-CoV-2 pseudovirus. The HuNAbs with high neutralizations were color-coded.

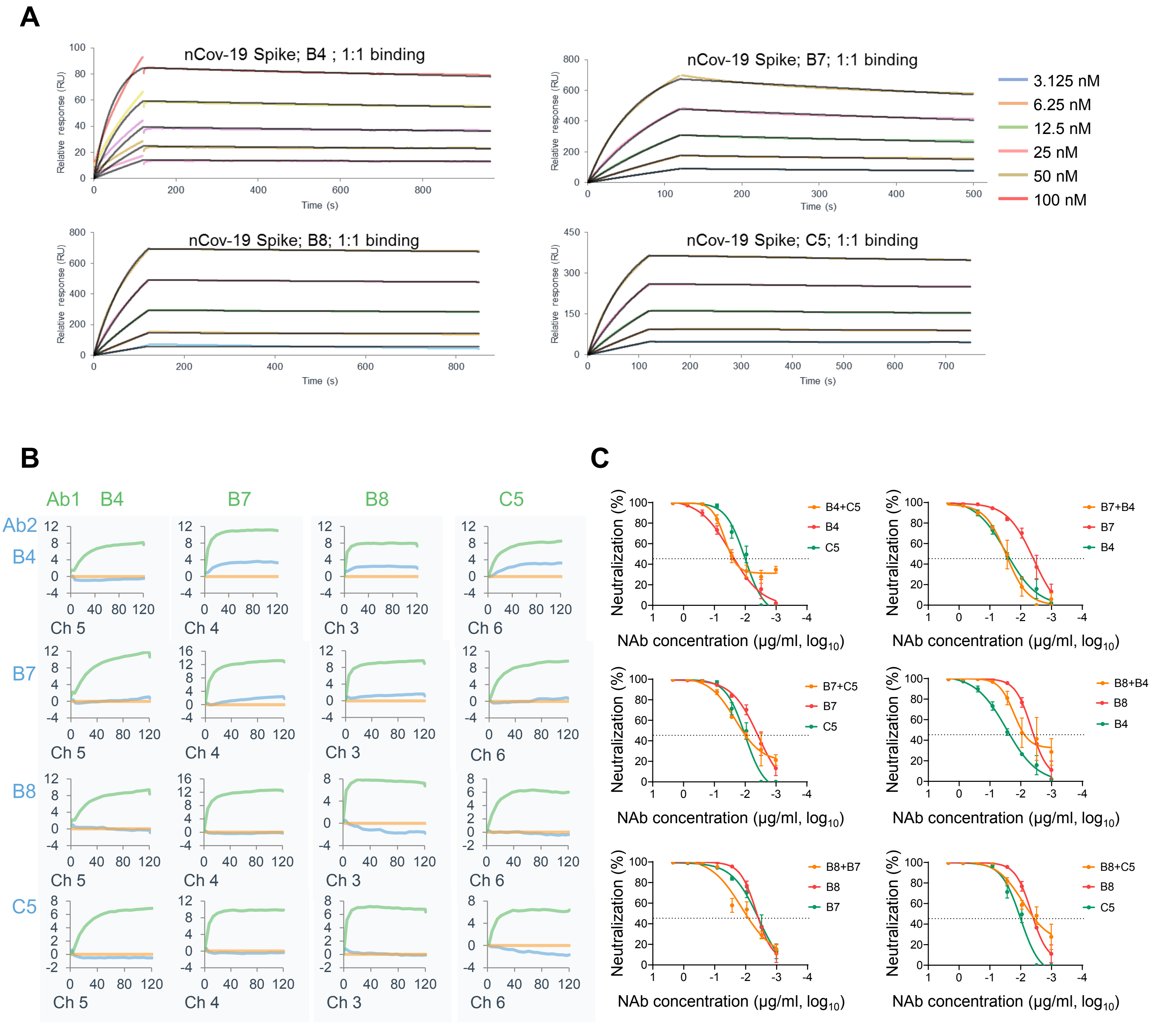

**Supplementary Fig. 2. Characterization of purified HuNAbs *in vitro*.**

**(A)** The binding dynamics of four most potent HuNAbs (B4, B7, B8 and C4) to SARS-CoV-2 Spike glycoprotein.

**(B)** The competitive binding between these four HuNAbs to SARS-CoV-2 RBD. Orange curve: the baseline; Green curve: the binding of test antibody to RBD; Blue Curve: the binding of test antibody (Ab1) to RBD after pre-incubation with the competitor antibody (Ab2).

**(C)** Lack of synergistic effect between pairs of these four HuNAbs by neutralization assay against the SARS-CoV-2 pseudovirus. The combined antibodies were mixed at 1:1 ratio.

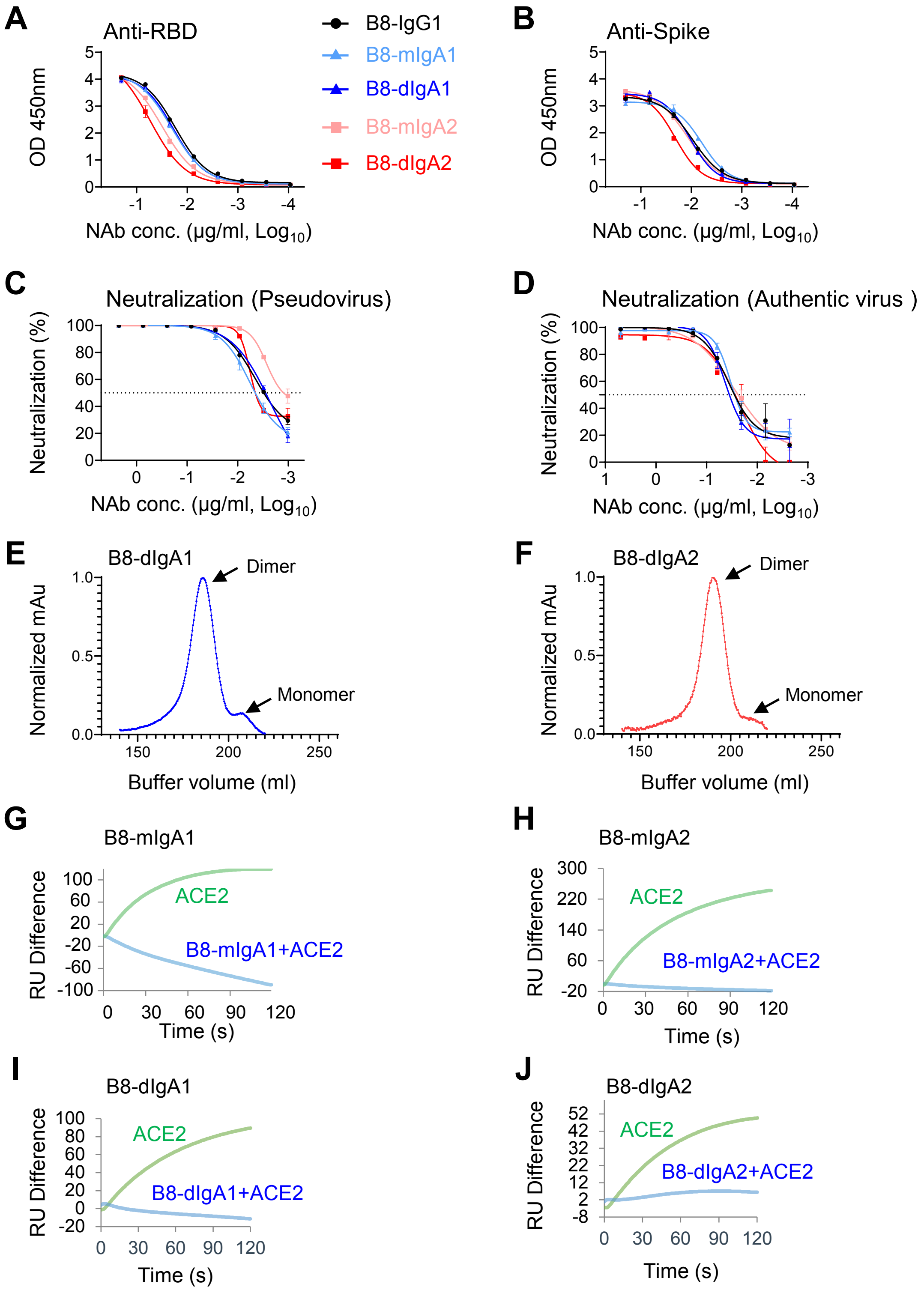

**Supplementary Fig. 3. Characterization of B8-based IgA NAbs.**

**(A)** RBD-specific binding activities of B8-mIgA1, B8-mIgA1, B8-dIgA1 and B8-dIgA2 as compared to B8-IgG1 measured by ELISA. **(B)** Spike-specific binding activities of B8-mIgA1, B8-mIgA1, B8-dIgA1 and B8-dIgA2 as compared to B8-IgG1 measured by ELISA. **(C)** Neutralization activities of B8-mIgA1, B8-mIgA1, B8-dIgA1 and B8-dIgA2 as compared to B8-IgG1 measured by decreased pseudotyped SARS-CoV-2 infection in HEK 293T-ACE2 cells. **(D)** Neutralization activities of B8-mIgA1, B8-mIgA1, B8-dIgA1 and B8-dIgA2 as compared to B8-IgG1 measured by decreased authentic SARS-CoV-2 infection in Vero-E6 cells. All the assays above (A-D) were performed in duplicates and the mean of the duplicates was shown with SEM. The antibody concentration in the x-axis is shown in log-transformed units. **(E)** The purity of dimeric B8-dgA1 was confirmed by size exclusion chromatography (SEC). **(F)** The purity of dimeric B8-dgA2 was confirmed by SEC. **(G-J)** The curves show binding of ACE2 to SARS-CoV-2 RBD with (blue) or without (blue) pre-incubation with B8-mIgA1, B8-mIgA2, B8-dIgA1, and B8-dIgA2, respectively, as measured by SPR.

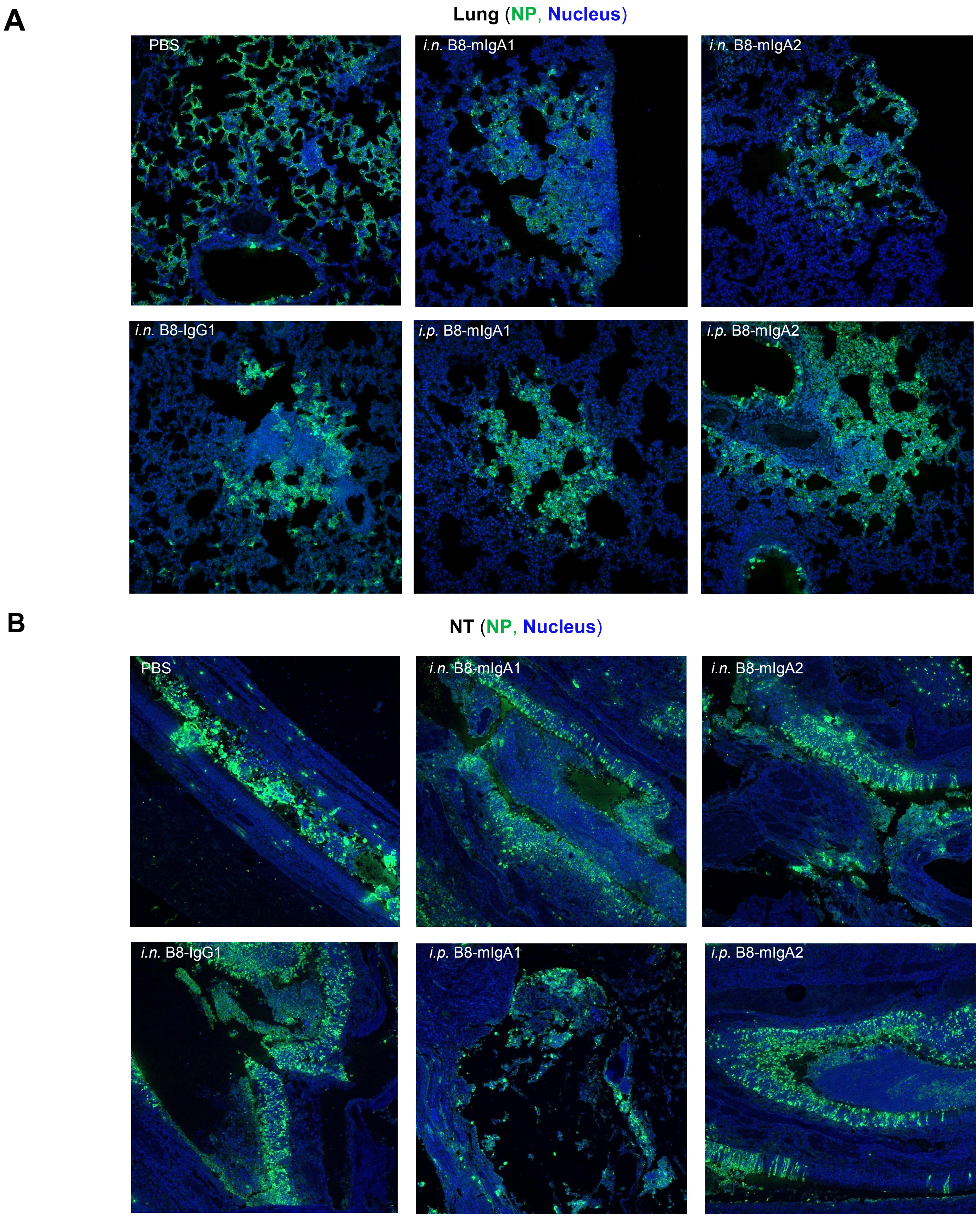

**Supplementary Fig. 4. SARS-CoV-2 infection at 4 dpi in both lung and NT of infected Syrian hamsters pre-treated with B8-mIgA1 or B8-mIgA2 by confocal microscope.**

**(A-B)** Representative images (100×) of infected foci in lungs **(A)** and NT **(B)** from each group as determined by anti-NP immunofluorescence (IF) staining. The SARS-CoV-2 NP and cell nuclei were stained with rabbit anti-SARS-CoV-2 NP (green) and DAPI (blue), respectively.

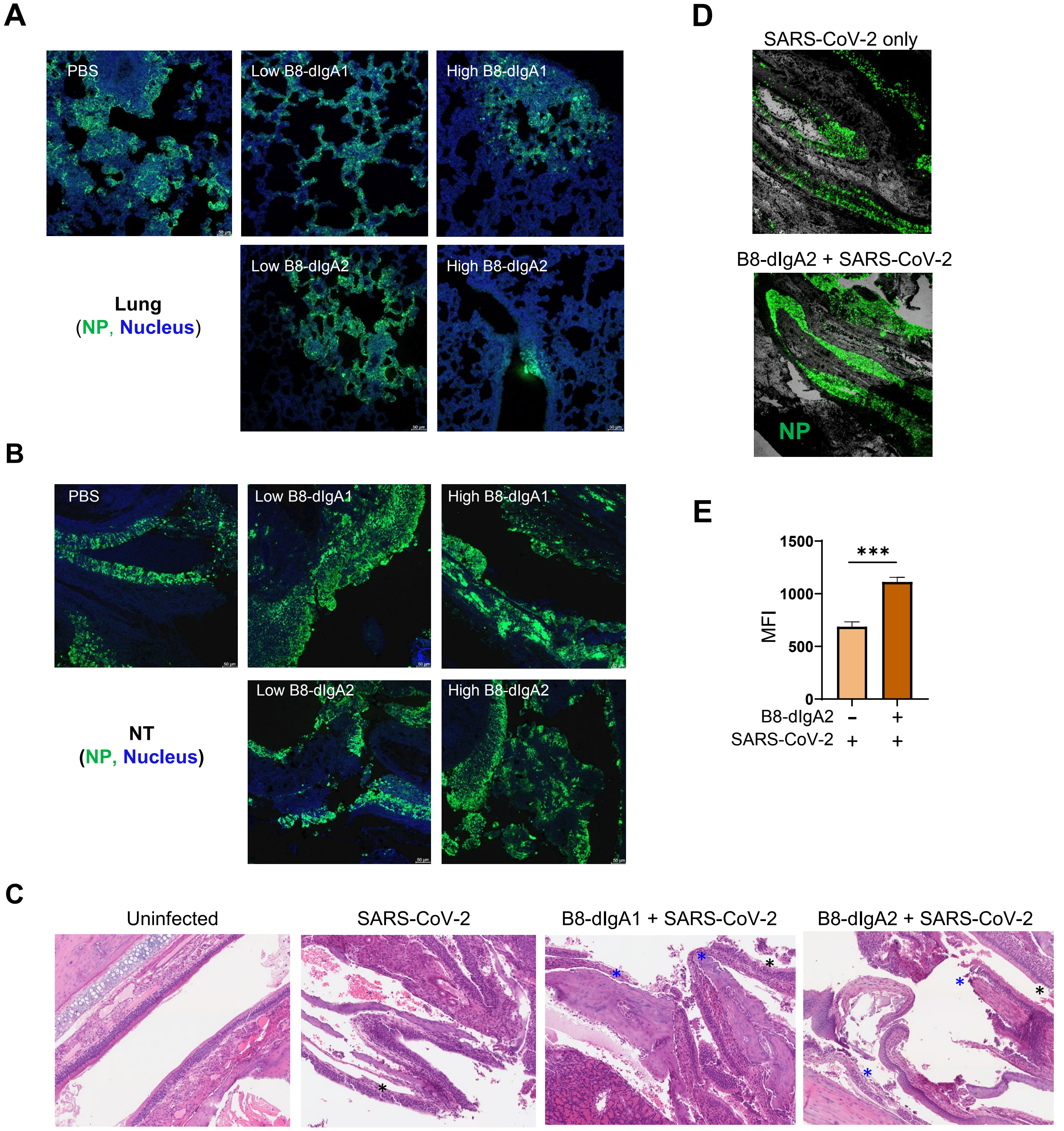

**Supplementary Fig. 5. SARS-CoV-2 infection at 4 dpi in both lung and NT of infected Syrian hamsters pre-treated with B8-dIgA1 or B8-dIgA2 by confocal and HE microscope.**
**(A-B)** Representative images (100×) of infected foci in lungs **(A)** and NT **(B)** from each group as determined by anti-NP immunofluorescence (IF) staining. The SARS-CoV-2 NP and cell nuclei were stained with rabbit anti-SARS-CoV-2 NP (green) and DAPI (blue), respectively.
**(C)** SARS-CoV-2 infection results in more extensive and severe damage of the NT epithelium in B8-dIgA-administrated animals (10× HE images). Injury of the NT epithelium is extensive with partial (black asterisk) or complete (blue asterisk) desquamation in B8-dIgA1- and B8-dIgA2-pretreated Syrian hamsters.
(**D**) Density of NP-positive cells in NT epithelium during live SARS-CoV-2 infection (50× images).
**(E)** Analysis of NP-positive MFI in 10 random NP-positive areas. Statistics were generated using the student *t* test. *$p$<0.05; **$p$<0.01; ***$p$<0.001.

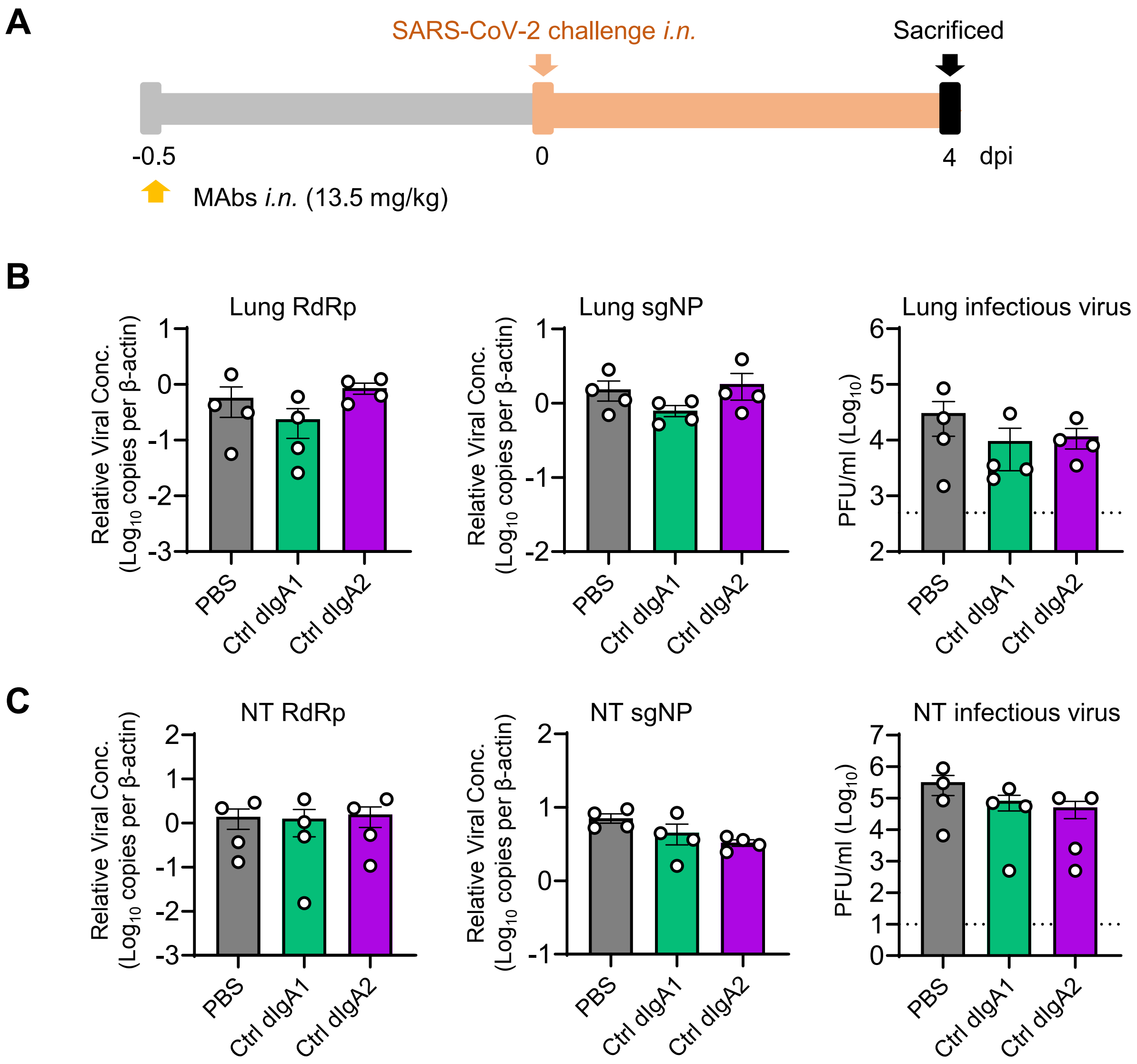

**Supplementary Fig. 6. Control dIgA1 or dIgA2 did not mediate enhancement of SARS-CoV-2 infection in NT.**

**(A)** Experiment schedule. Two groups of hamsters (n=4 per group) were inoculated intranasally with control dIgA1 and control dIgA2 at a high dose of 13.5 mg/kg 12 hours before intranasal viral challenge, respectively. Another group of hamsters (n=4) received PBS as control. On day 0, each hamster was intranasally challenged with a dose of $10^5$ PFU of SARS-CoV-2 as mentioned in Figure 5A. All hamsters were sacrificed on 4 dpi for analysis.

**(B)** The viral loads in lung were determined by three assays.

**(C)** The viral loads in NT were determined by three assays.

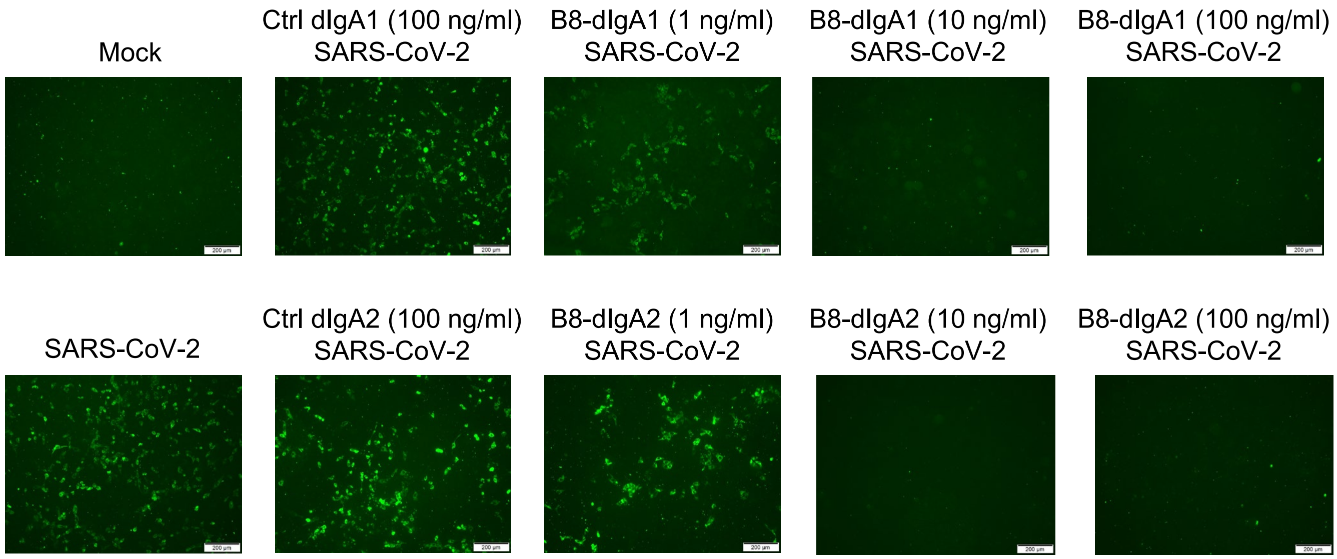

**Supplementary Fig. 7. Potent neutralization of live SARS-CoV-2 infection by B8-dIgA1 and B8-IgA2 in human kidney cell line HK-2.**

The IF staining of SARS-CoV-2 NP (in green) in infected HK-2 cells pre-treated with different dose of antibody as indicated. The representative image of each group was shown. Scale bars represent 200 μm.

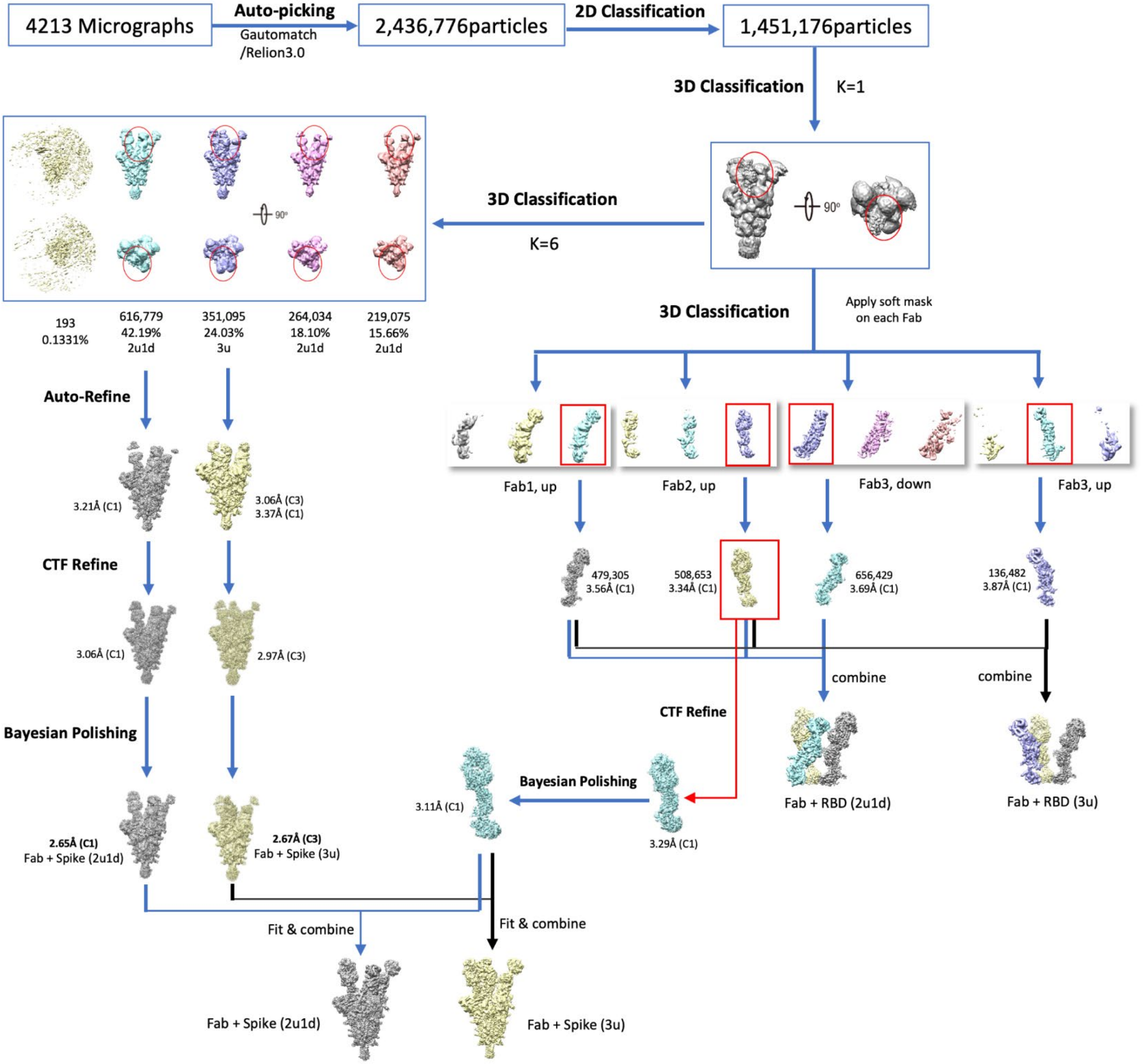

**Supplementary Fig. 8. A flow-chart of SARS-CoV-2 S-B8 complex cryo-EM data processing.** Different map density of RBD-Fab portions were emphasized in red cycle.

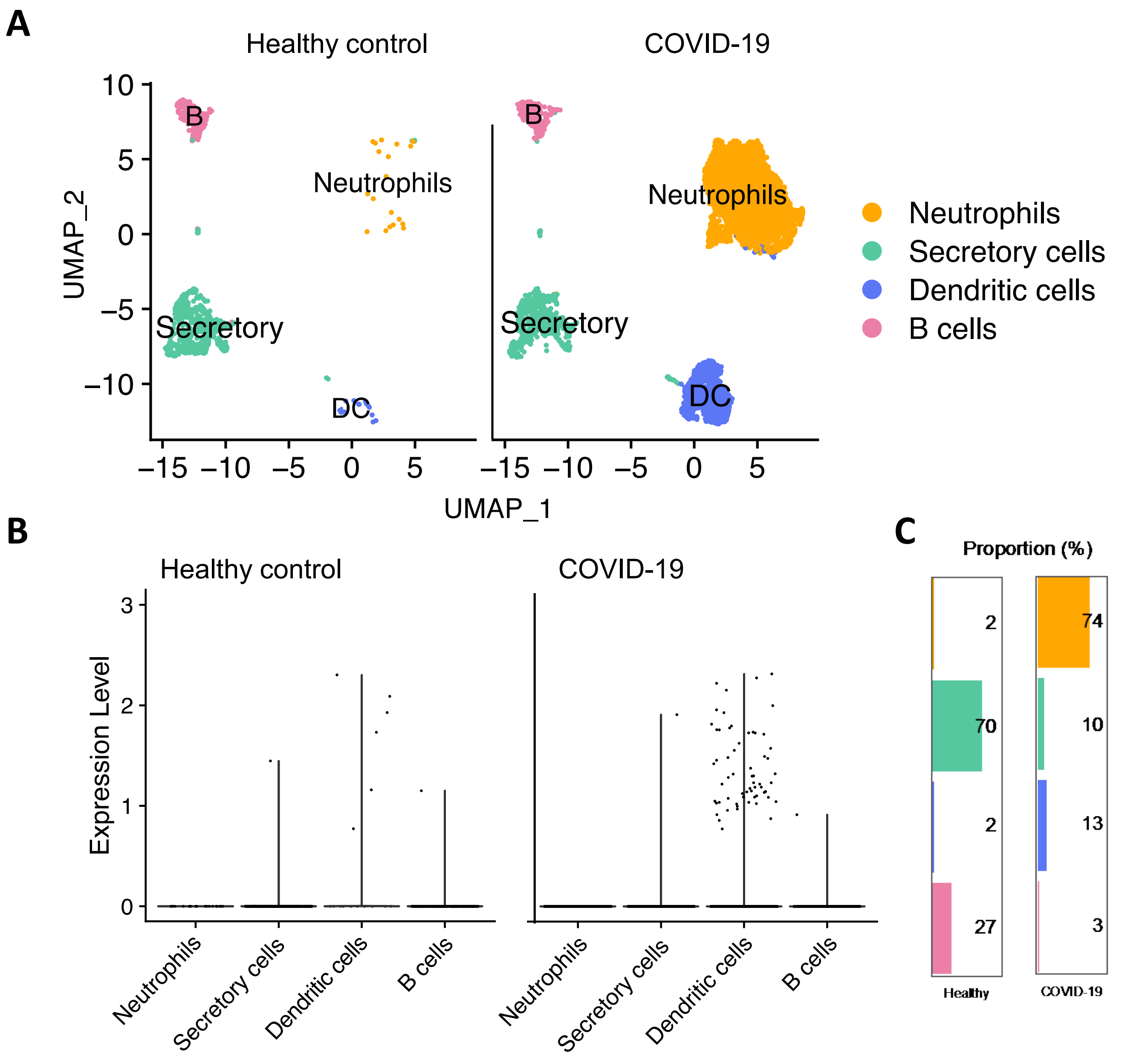

**Supplementary Fig. 9. Preliminary analysis of the human nasal cytology data.**
**(A)** The analysis was based data submitted under accession code GSE171488 (healthy donor nasal brushing) and GSE164547 (COVID-19 patient nasal brushing). Among CD14 positive cells, the neutrophil and monocyte-differentiated DCs were mainly increased in the nasal cytology samples of COVID-19 patients.
**(B)** Increased CD209 expression on nasal DCs of COVID-19 patients.
**(C)** The proportion of nasal DCs increased 6.5-fold from 2% to 13% in nasal samples compared between health and COVID-19 subjects.

Supplementary Table 1. Clinical characteristics of SARS-CoV-2 infected subjects.

| Patient ID | Age | Gender | Days after symptom onset (or 1st hospitalization) | Disease Severity |
|---|---|---|---|---|
| P1 | 56 | F | 114 | Severe |
| P2 | 62 | M | 116 | Severe |
| P3 | 21 | M | 49 | Asymptomatic |
| P4 | 75 | M | 15 | Mild |

Supplementary Table 2. Gene family analysis of four HuNAbs.

| HuNAbs | Heavy chain | | | | Light chain | | | |
|---|---|---|---|---|---|---|---|---|
| | IGHV | IGHJ | CDR3 length | SHM (%) | IGKV | IGKJ | CDR3 length | SHM (%) |
| B4 | IGHV3-66*01, IGHV3-66*04 | IGHJ6*02 | 12 | 3.8 | IGKV1-33*01, IGKV1D-33*01 | IGKJ5*01 | 9 | 4.6 |
| B7 | IGHV1-69*18 | IGHJ6*02 | 18 | 0.0 | IGKV3-11*01 | IGKJ2*01 | 9 | 0.7 |
| B8 | IGHV1-69*18 | IGHJ6*02 | 14 | 4.8 | IGKV3-11*01 | IGKJ4*01 | 9 | 1.7 |
| C5 | IGHV1-69*18 | IGHJ4*02 | 16 | 2.4 | IGKV3-20*01 | IGKJ1*01 | 9 | 2.8 |

Supplementary Table 3. Binding ability of HuNAbs to SARS-CoV-2 RBD and spike.

| HuNAbs | SARS-CoV-2 RBD | SARS-CoV-2 spike |
|---|---|---|
| | $EC_{50}$ (μg/ml) | $EC_{50}$ (μg/ml) |
| A6 | 0.30 | 17.94 |
| B4 | 0.06 | 0.06 |
| B7 | 0.04 | 0.018 |
| B8 | 0.02 | 0.02 |
| C5 | 0.03 | 0.03 |

Supplementary Table 4. Neutralization capability of RBD-specific HuNAbs.

| HuNAbs | Pseudovirus | | Live virus | |
|---|---|---|---|---|
| | $IC_{50}$ (μg/ml) | $IC_{90}$ (μg/ml) | $IC_{50}$ (μg/ml) | $IC_{90}$ (μg/ml) |
| B4 | 0.029 | 0.136 | 0.048 | 0.134 |
| B7 | 0.015 | 0.094 | 0.030 | 0.060 |
| B8 | 0.0095 | 0.046 | 0.013 | 0.032 |
| C5 | 0.038 | 0.083 | 0.024 | 0.044 |

Supplementary Table 5. Surface plasmon resonance analysis of RBD-specific HuNAbs.

| Antigen | Analyte | Kinetics model | ka (1/Ms) | kd (1/s) | KD (M) | tc |
|---|---|---|---|---|---|---|
| Spike | B4 | 1:1 | 2.11e+05 | 9.85e-05 | 4.66e-10 | 1.32e+13 |
| Spike | B7 | 1:1 | 3.28e+05 | 4.35e-04 | 1.32e-09 | 2.08e+09 |
| Spike | B8 | 1:1 | 2.24e+05 | 3.78e-05 | 1.69e-10 | 1.42e+12 |
| Spike | C5 | 1:1 | 2.50e+05 | 7.70e-05 | 3.08e-10 | 1.56e+09 |

Supplementary Table 6. B8-IgG1 concentrations in different compartments of hamsters.

| Group | Inoculation time (hour) | Inoculation dose (mg/Kg) | Day 0 | Day 4 | | |
|---|---|---|---|---|---|---|
| | | | Serum (ng/ml) | Serum (ng/ml) | Lung homogenate (ng/ml) | Nasal turbinate homogenate (ng/ml) |
| G1 | -24 | *i.p.* 1.5 | 4257 ± 1517 | 2101 ± 1039 | 128.5 ± 13.18 | 20.66 ± 17.53 |
| G2 | 24 | *i.p.* 1.5 | *N.A.* | 4868 ± 797.8 | 238.1 ± 28.68 | 86.20 ± 18.73 |
| G3 | 48 | *i.p.* 1.5 | *N.A.* | 4135 ± 1674 | 229.3 ± 47.89 | 93.82 ± 8.302 |
| G4 | 72 | *i.p.* 1.5 | *N.A.* | 3252 ± 1749 | 192.4 ± 43.07 | 46.01 ± 3.934 |

*N.A.*: Not applicable

Supplementary Table 7. Binding of B8 HuNAbs to SARS-CoV-2 RBD and spike.

| HuNAbs | SARS-CoV-2 RBD | SARS-CoV-2 Spike |
|---|---|---|
| | $EC_{50}$ (μg/ml) | $EC_{50}$ (μg/ml) |
| B8-IgG1 | 0.02 | 0.02 |
| B8-mIgA1 | 0.020 | 0.006 |
| B8-mIgA2 | 0.031 | 0.012 |
| B8-dIgA1 | 0.019 | 0.011 |
| B8-dIgA2 | 0.054 | 0.021 |

Supplementary Table 8. Neutralization potency of B8 HuNAbs.

| HuNAbs | Pseudovirus | | Live virus | |
|---|---|---|---|---|
| | $IC_{50}$ (μg/ml) | $IC_{90}$ (μg/ml) | $IC_{50}$ (μg/ml) | $IC_{90}$ (μg/ml) |
| B8-IgG1 | 0.0095 | 0.046 | 0.013 | 0.032 |
| B8-mIgA1 | 0.012 | 0.052 | 0.012 | 0.018 |
| B8-mIgA2 | 0.0057 | 0.014 | 0.010 | 0.051 |
| B8-dIgA1 | 0.0048 | 0.033 | 0.015 | 0.026 |
| B8-dIgA2 | 0.012 | 0.021 | 0.008 | 0.026 |

Supplementary Table 9. HuNAb concentration in different tissue compartments of infected hamsters.

| HuNAbs | Inoculation time (hour) | Inoculation dose (mg/Kg) | Day 0 | Day 4 | | |
|---|---|---|---|---|---|---|
| | | | Serum (ng/ml) | Serum (ng/ml) | Lung homogenate (ng/ml) | Nasal turbinate homogenate (ng/ml) |
| B8-mIgA1 | -24 | *i.n.* 4.5 | 90.71 ± 11.60 | *U.D.* | 16.54 ± 7.178 | *U.D.* |
| | -24 | *i.p.* 4.5 | 8682 ± 1749 | *U.D.* | *U.D.* | *U.D.* |
| B8-mIgA2 | -24 | *i.n.* 4.5 | 21.36 ± 6.415 | *U.D.* | *U.D.* | *U.D.* |
| | -24 | *i.p.* 4.5 | 1561 ± 129.1 | *U.D.* | *U.D.* | *U.D.* |
| B8-dIgA1 | -12 | *i.n.* 4.5 | 3.784 ± 2.081 | *U.D.* | 85.99 ± 17.52 | *U.D.* |
| | -12 | *i.n.* 13.5 | 1.981 ± 0.2688 | *U.D.* | 88.48 ± 23.44 | *U.D.* |
| B8-dIgA2 | -12 | *i.n.* 4.5 | *U.D.* | *U.D.* | 6.383 ± 4.624 | *U.D.* |
| | -12 | *i.n.* 13.5 | *U.D.* | *U.D.* | 409.1 ± 108.0 | *U.D.* |

*U.D.*: undetectable

Supplementary Table 10. HuNAb concentration in different tissue compartments of naive Syrian hamsters at the time of viral challenge.

| HuNAbs | Inoculation time (hour) | Inoculation dose (mg/Kg) | Day 0 | | | |
|---|---|---|---|---|---|---|
| | | | Nasal wash (ng/ml) | Serum (ng/ml) | Lung homogenate (ng/ml) | Nasal turbinate homogenate (ng/ml) |
| B8-IgG1 | -12 | *i.n.* 4.5 | 208.6 ± 23.23 | 479.9 ± 62.53 | 24336 ± 2661 | 121.1 ± 26.38 |
| B8-mIgA1 | -12 | *i.n.* 4.5 | 315.9 ± 186.1 | 18.84 ± 5.802 | 18914 ± 1672 | 53.87 ± 11.14 |
| B8-mIgA2 | -12 | *i.n.* 4.5 | 204.1 ± 25.18 | 16.05 ± 2.515 | 32712 ± 12440 | 116.5 ± 72.62 |
| B8-dIgA1 | -12 | *i.n.* 4.5 | 43.18 ± 27.52 | 6.827 ± 6.520 | 4347 ± 1598 | 14.74 ± 5.448 |
| B8-dIgA2 | -12 | i.n. 4.5 | 56.79 ± 15.69 | 19.00 ± 7.562 | 28033 ± 12575 | 23.74 ± 1.445 |

Supplementary Table 11. Statistics of cryo-EM data collection, processing and model refinement.

| Data collection | | | | | | |
|---|---|---|---|---|---|---|
| EM equipment | Titan Krios | | | | | |
| Voltage (kV) | 300 | | | | | |
| Detector | K3 Summit | | | | | |
| Electron dose ($e^{-}/Å^{2}$) | 50 | | | | | |
| Defocus range (μm) | -1.0 ~ -2.0 | | | | | |
| Pixel size (Å) | 1.0979 | | | | | |
| Reconstruction | | | | | | |
| Software | Relion 3.0 & Relion 3.1 | | | | | |
| Structure & state | Spike | | RBD-Fab | | | |
| | 3u | 2u1d | #1, up | #2, up | #3,down | #3,up |
| Symmetry imposed | C3 | C1 | C1 | C1 | C1 | C1 |
| Final particles | 616,779 | 351,095 | 479,305 | 508,653 | 656,429 | 136,482 |
| Map resolution (Å) | 2.76 | 2.65 | 3.56 | 3.11 | 3.69 | 3.87 |
| Atomic modeling | | | | | | |
| Software | Chimera, Coot, Phenix | | | | | |
| R.m.s. deviations | | | | | | |
| Bond lengths (Å) | 0.003 | 0.004 | 0.004 | 0.003 | 0.003 | 0.03 |
| Bond angles (°) | 0.591 | 0.601 | 0.815 | 0.683 | 0.628 | 0.794 |
| MolProbity score | 1.63 | 1.72 | 2.05 | 1.86 | 2.09 | 2.29 |
| Clash score | 5.36 | 6.32 | 9.99 | 7.21 | 12.05 | 17.48 |
| Poor rotamers (%) | 0 | 0 | 0 | 0 | 0.19 | 0 |
| Ramachandran plot (%) | | | | | | |
| Favored | 95.15 | 94.55 | 90.66 | 92.5 | 91.69 | 90.24 |
| Allowed | 4.85 | 5.45 | 9.34 | 7.5 | 8.31 | 9.76 |
| Outliers | 0 | 0 | 0 | 0 | 0 | 0 |

Supplementary Table 12. Contacts between SARS-CoV-2 RBD and B8 Fab (distance cutoff 4Å).

| RBD | Heavy chain | RBD | Light chain |
|---|---|---|---|
| V445 | E1 | E484 | W94 |
| G446* | K98 | G485 | W94 |
| Y449* | N31, Y32, K98 | F486* | N93, W94 |
| N450 | N31 | N487* | N93 |
| L452 | N31, F55, L101 | Y489* | S92, N93 |
| F456* | L104 | T500* | T56 |
| T470 | F55 | | |
| I472 | T57 | | |
| V483 | N59 | | |
| E484 | R50, L52, T57, N59, F103 | | |
| C488 | F103 | | |
| Y489* | F103, L104 | | |
| F490 | F55, L101 | | |
| L492 | L101, A102 | | |
| Q493* | A102, L104 | | |
| S494 | N31, L101 | | |

* ACE2 binding sites

Supplementary Table 13. Neutralization of SARS-CoV-2 variants by B8-derived HuNAbs.

| Fold change of IC50 from WT (D614G) | | RBD-specific HuNAb | | | | |
|---|---|---|---|---|---|---|
| | | B8-IgG1 | B8-mIgA1 | B8-dIgA1 | B8-mIgA2 | B8-dIgA2 |
| UK (B.1.1.7) | UKΔ8 | -30.4 | -25.3 | -6.5 | -19 | -22.5 |
| | 69-70del | -4.5 | -0.2 | -0.3 | 0.2 | -1.5 |
| | N501Y | -3.7 | -3 | -5 | -2.3 | -3.9 |
| | A570D | 0.4 | 0.6 | 0.5 | -0.4 | -0.5 |
| | P681H | -0.1 | 0.7 | 0.5 | 0.5 | 0.6 |
| | S982A | 0.6 | 0.4 | 0.2 | 0.4 | 0 |
| | D1118H | -4.8 | -3.2 | -2.3 | -2.4 | -3 |
| SA (B.1.351) | SAΔ9 | -71.7 | <-1000 | <-1000 | -81.8 | -549 |
| | L18F | -5.6 | -0.1 | -0.1 | -1.1 | -0.9 |
| | D80A | -4.5 | -3.3 | -1.3 | -0.3 | -0.9 |
| | D215G | -22 | -26.4 | -11.5 | -21.3 | -15.3 |
| | 242-244 del | -5.2 | -0.5 | -3.2 | -0.1 | -0.9 |
| | R246I | -2.4 | -0.7 | 0.3 | 0.3 | -0.4 |
| | K417N | -1.7 | 0.5 | 0.3 | -0.2 | -1.9 |
| | E484K | -779.5 | <-1000 | <-1000 | <-1000 | <-1000 |
| | N501Y | -3.7 | -3 | -5 | -2.3 | -3.9 |
| | A701V | -0.1 | -0.4 | 0.3 | 0.7 | 0.4 |
| Others | G485S | -1.3 | -1.8 | -2.1 | 0.7 | -0.6 |
| | F486A | -5.6 | 0.9 | -1 | 0.5 | -0.9 |
| | T716I | -1.3 | -3.5 | -0.9 | -13.1 | -5 |
| | Y453F | -0.6 | -5.3 | -2.3 | -0.5 | 0.8 |

Note: red, resistant; green, sensitive
